# Profiling novel lateral gene transfer events in the human microbiome

**DOI:** 10.1101/2023.08.08.552500

**Authors:** Tiffany Y. Hsu, Etienne Nzabarushimana, Dennis Wong, Chengwei Luo, Robert G. Beiko, Morgan Langille, Curtis Huttenhower, Long H. Nguyen, Eric A. Franzosa

**Affiliations:** Harvard T.H. Chan School of Public Health, Boston, MA, USA; Massachusetts General Hospital and Harvard Medical School, Boston, MA, USA; Faculty of Computer Science, Dalhousie University, Halifax, Nova Scotia, Canada; The Broad Institute of MIT and Harvard, Cambridge, MA, USA; Department of Pharmacology, Dalhousie University, Halifax, Nova Scotia, Canada

**Keywords:** Horizontal gene transfer, human microbiome, metagenomics

## Abstract

Lateral gene transfer (LGT) is an important mechanism for genome diversification in microbial populations, including the human microbiome. While prior work has surveyed LGT events in human-associated microbial isolate genomes, the scope and dynamics of novel LGT events arising in personal microbiomes are not well understood, as there are no widely adopted computational methods to detect, quantify, and characterize LGT from complex microbial communities. We addressed this by developing, benchmarking, and experimentally validating a computational method (WAAFLE) to profile novel LGT events from assembled metagenomes. Applying WAAFLE to >2K human metagenomes from diverse body sites, we identified >100K putative high-confidence but previously uncharacterized LGT events (∼2 per assembled microbial genome-equivalent). These events were enriched for mobile elements (as expected), as well as restriction-modification and transport functions typically associated with the destruction of foreign DNA. LGT frequency was quantifiably influenced by biogeography, the phylogenetic similarity of the involved taxa, and the ecological abundance of the donor taxon. These forces manifest as LGT networks in which hub species abundant in a community type donate unequally with their close phylogenetic neighbors. Our findings suggest that LGT may be a more ubiquitous process in the human microbiome than previously described. The open-source WAAFLE implementation, documentation, and data from this work are available at http://huttenhower.sph.harvard.edu/waafle.

## Introduction

Lateral gene transfer (LGT), or the movement of genetic material between organisms through means other than vertical inheritance from parent to offspring, is a major force in the evolution and diversification of microbes^1–6^. Indeed, studies estimate that 10-20% of genes in some bacterial clades were acquired by LGT^7–9^. Also referred to as horizontal gene transfer (HGT), LGT is thought to be an important process in microbial communities, as it may provide recipients with advantageous traits, such as antibiotic resistance, the ability to degrade certain compounds, or survive in different environmental niches^10^. However, the majority of methods for detecting LGT have focused on isolate genomes, making it difficult to assess the prevalence, modes, and functions of LGT in complex microbial communities.

Prior studies of LGT in the human microbiome have applied traditional methods of LGT event detection to isolate genomes, typically using phylogenetic approaches based on gene-species tree construction and reconciliation or relying on composition-based inference to identify candidate acquisitions^11–16^. Such studies have revealed enrichments for LGT among microbes native to oral and gut sites and that LGT rates were influenced by host^17,18^ (e.g. lifestyle and geography) and microbial attributes (e.g. phylogeny and ecology)^19^. For example, comparisons of gut microbiomes from Fijian and North American individuals revealed differences in transferred genes linked to diet and geography^20^. Another study found that LGT events occur more frequently in industrialized and urban gut microbial communities^17^. Moreover, while LGT is more common between phylogenetically related species^21,22^, this trend was secondary to LGT enrichment among species associated with specific environments, such as the human body^11^. A recent study also found that LGT from maternal gut bacteria may drive infant development through the sharing of functions associated with immunity and dietary changes^10^. These findings are consistent with a proposed theory that LGT is highly adaptive among niche-sharing microbes facing dynamic environmental pressures^17,18,23^, an idea further supported by observed LGT-induced enrichments for survival-related pathogenicity factors like antimicrobial-resistance genes and carbohydrate usage pathways^11,24,25^. However, while LGT acts as an ecological force where acquisition of adaptive genes may benefit recipients and promote community stability^19^, LGT could also negatively impact the host and its microbiome^18^, underscoring the need to rigorously study this phenomenon.

While studying microbial community LGT from isolate genomes has thus avoided the challenges of culture-independent methodologies, the strategy suffers from several drawbacks. Foremost is that the set of LGT events involving a recipient species must be gleaned from one or a few reference genomes per organism. Therefore, variation in the occurrence and fixation of LGT events within species may go undetected, resulting in a dramatic underestimation of LGT-based strain personalization. Applying these conventional LGT detection methods on metagenome assemblies may also have practical limitations due to the fact that LGT contigs do not bin well, potentially as a consequence of complex flanking repeat regions that can result in loss of coverage^2^. Additionally, assessing LGT from a single reference genome obscures its evolutionary history within environments. For example, evolutionary trajectories vary for human-associated microbes from ancient (i.e. predating the origin of modern humans) to those arising within the host’s lifetime (herein referred to as “recent” LGT events). While such limitations could in principle be ameliorated by assembly of complete microbial genomes, this process is computationally challenging and limited to the highest-coverage species, particularly when analyzing difficult-to-assemble mobile elements.

Relatively few methods have been specifically designed to identify and profile LGT events in microbial communities. Notably, general methods of LGT detection based on variability in sequence composition (e.g. Alien_Hunter^26^ and DarkHorse^27^) can theoretically be applied to metagenomic contigs and/or bins, though their effectiveness in that context is unclear. Daisy^28^ was among the first methods to specifically target LGT events in microbial communities, which it accomplished by mapping metagenomic reads to putative donor and recipient genomes. The need to pre-specify these genomes limits the method’s utility in comprehensive community LGT profiling, though the more recent DaisyGPS^29^ has been developed to aid in donor and recipient selection. Other methods, such as LEMON^30^, follow a related approach of comparing reference genomes with metagenomic reads to identify potential LGT breakpoints in the underlying community strains. In contrast, MetaCHIP^31^ identifies LGT events between microbial community members by inspecting their metagenome-assembled genomes (MAGs) for discordance between species and gene trees (independent of external reference genomes). While this design makes MetaCHIP a highly general community LGT profiler, it is expected to lack sensitivity to LGT events occurring outside of a sample’s higher-quality MAGs or involving genetic material from outside the community. Thus, a method that can profile microbial community LGT both broadly and accurately remains an unmet need.

To address these issues, we developed a phylogenetically agnostic computational method for novel LGT detection and profiling from shotgun metagenomic assemblies which we call WAAFLE (**W**orkflow to **A**nnotate **A**ssemblies and **F**ind **L**GT **E**vents). We benchmarked WAAFLE on highly fragmented synthetic assemblies, identifying the majority of expected spiked-in LGT with <0.5% false-positive detection rate and with improved sensitivity compared to existing community-applicable methods. We then carried out the first comprehensive culture-independent profiling of LGT across diverse human body sites, drawing on >2,000 assembled metagenomes from 264 individuals and 16 body sites from the expanded Human Microbiome Project (HMP1-II)^32^. We identified over 100,000 high-confidence novel LGT events (with “novel” defined here as “not previously observed in microbial isolate genomes”). We also experimentally validated high-confidence candidates in a second independent metagenomic cohort. WAAFLE was thus used to interrogate a wide variety of complex microbial communities, and these results considerably expand our understanding of the network of transferring species, functions, and general determinants of LGT among human-associated microbes.

## Results

### Identifying novel LGT from metagenome assemblies

As input, WAAFLE uses assembled metagenomes as a collection of contigs (**Fig. 1A**). The method is robust to assemblies that are neither binned nor particularly complete. WAAFLE compares each contig to a taxonomically annotated reference database of microbial gene sequences using a homology-based search. The contig’s protein-coding open reading frames (ORFs), provided as input or identified during search, are then analyzed using an iterative, two-step taxonomic placement process. WAAFLE first determines whether the contig’s complete set of ORFs can be reasonably explained by a single species. If so, WAAFLE assigns the contig to that species. If not, WAAFLE then determines whether any two species can jointly explain the ORFs with confidence. If so, WAAFLE proposes a putative LGT between those species. If not, WAAFLE repeats this process at the next taxonomic rank (i.e. genus, family, and so forth). In each case, to “explain” a contig, each of the contigs’ ORFs must align to one or two clades’ pangenomes above a prespecified homology threshold (defined as k_1_ for single-clade explanations and k_2_ for clade-pair/LGT explanations, respectively, **Methods** and **Fig. S1**).

**Figure 1.**
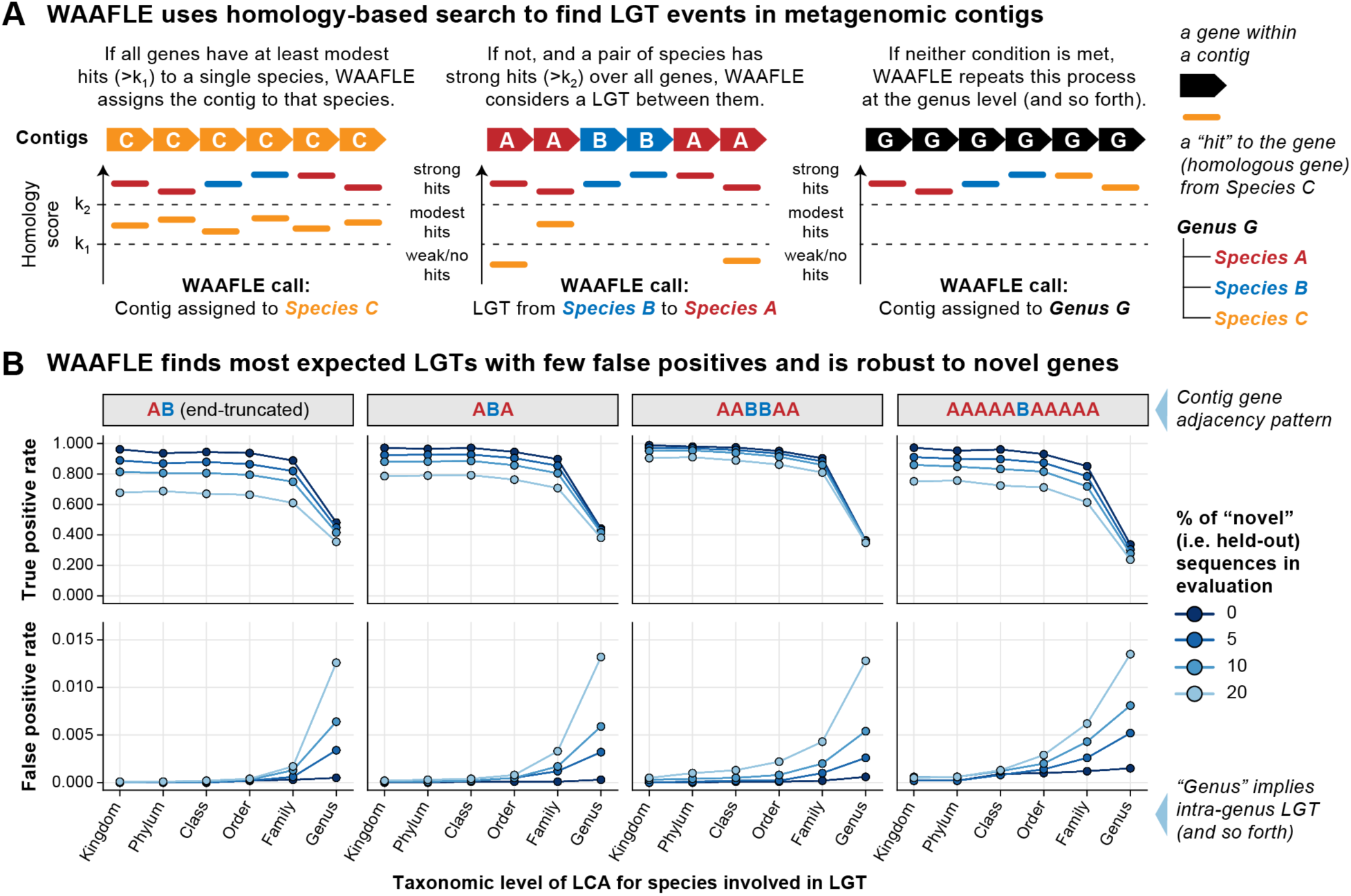
The WAAFLE algorithm is accurate for novel lateral transfer discovery in metagenomes. (**A**) WAAFLE identifies putative LGT events in metagenomes by aligning metagenomic contigs to microbial reference sequences in order to i) taxonomically place the contigs and ii) identify potential LGT events contained within them, iteratively from intra-genus to more distant transfer events. (**B**) We evaluated WAAFLE on synthetic contigs of different synteny (gene order) configurations assembled from recipient (‘A’) and donor (‘B’) genomes. In the top row (true positive rate calculation), taxonomic level indicates the LCA of the donor and recipient species involved in an LGT-containing contig. In the bottom row (false positive rate calculation), taxonomic level stratifies erroneous LGT calls according to remoteness. To determine LGT directionality, if a gene adjacency pattern matches A+B+A+ in a contig passing quality control (**Methods**), then A is considered the recipient and B the donor.

### WAAFLE identifies LGT events with high specificity

We optimized WAAFLE’s parameters, evaluated its performance, and benchmarked it against existing methods using synthetic contigs with known LGT content (including control contigs with no LGT). To construct synthetic contigs, we selected pairs of isolate genomes (A and B) at a pre-specified taxonomic separation, defined here by the taxonomic rank of the genomes’ lowest common ancestor (LCA). We then randomly sampled, mutated, and concatenated genes from the two genomes to make LGT-containing contigs of a desired adjacency pattern (e.g. “ABA” corresponds to a transfer of a single B gene between two A genes). To generate control contigs without LGT, we performed the same procedure using pairs of isolate genomes from the same species. For each adjacency pattern and level of taxonomic separation, we generated 40 contigs from each of 250 genome pairs (for 10K total contigs) alongside equal numbers of control contigs (**Methods** and **Fig. S1**).

We first used these synthetic contigs to ensure that WAAFLE identified as many expected LGT events as possible while minimizing false-positive calls. Most critically, this involved selecting similarity thresholds for assigning a contig to a single clade vs. a pair of clades (the thresholds k_1_ and k_2_, respectively, **Fig. 1A**). Lower k_1_ and higher k_2_ make it easier to assign a contig to a single clade and more difficult to invoke LGT (**Fig. S2**). Although such settings reduce WAAFLE’s sensitivity, we favored them as defaults to avoid reporting weakly supported LGT events. Similarly, WAAFLE filters candidate LGT-containing contigs that could be explained by other biological mechanisms, such as gene deletion. For example, if a candidate LGT contig contains a large fraction of ambiguous ORFs or if the transferred gene is found in sister clades of the putative recipient, WAAFLE conservatively rejects the LGT (**Fig. S3** and **S4**). Lastly, we further improved accuracy by prefiltering low-confidence ORF calls prior to taxonomic placement (**Fig. S5**).

In an initial synthetic evaluation using default parameters, WAAFLE identified >84% of intergenus LGT events, i.e. those that occurred between taxa with an LCA above the genus level, and reported very few false positives (<0.1%, **Fig. 1B**). Intragenus LGT events, or those occurring within genera, were comparatively harder to detect (33-48% sensitivity) due to more extreme pangenome overlap between congeneric species (WAAFLE’s conservative approach invokes a single-species explanation for a contig before considering LGT of a shared gene, **Fig. 1A**). To evaluate WAAFLE’s accuracy in metagenomes for which well-matched pangenome references were not available, we repeated this evaluation while holding out 5 to 20% of reference sequences during WAAFLE’s homology-based search step. As expected, held-out sequences induced additional false positives, mostly at the intra-genus level and never exceeding 1.5% of control contigs. Relative to species divergence and database incompleteness, gene adjacency patterns, and partially assembled genes had minimal impact on WAAFLE’s ability to discriminate LGT and control contigs. Additionally, WAAFLE’s species-level taxonomic assignments were >97% accurate for LGT contigs and >90% accurate for control contigs, while genus-level assignments were >99% accurate (**Fig. S6**).

We next compared WAAFLE’s performance with that of more general sequence composition-based methods of LGT detection, specifically Alien_Hunter and DarkHorse, as applied to the synthetic LGT-containing and control contigs introduced above. While we understood that these methods were optimized for application to isolate genomes, we were surprised to observe the extent of their struggles with metagenomic contigs. Indeed, none of the synthetic contigs achieved the contiguous input sequence length required for analysis by Alien_Hunter, and it was thus excluded from further downstream quantitative evaluation. DarkHorse managed to produce interpretable outputs for a larger fraction of the contigs. However, it only produced a non-trivial fraction of true positive LGT calls when applied to the longest contigs (>10 genes) containing LGT between different phyla (41% sensitivity, **Fig. S7A**). In comparison, even when conservatively holding out 20% of its database, WAAFLE was 69-89% sensitive to remote LGT across a range of contig lengths and gene order configurations. WAAFLE additionally achieved a better worst-case false positive rate compared to DarkHorse (1.2% vs 1.5%, respectively).

We further compared WAAFLE to MetaCHIP, a method that uses homology-based search and phylogenetic validation to detect LGT from binned metagenome assemblies^31^. Given MetaCHIP’s expectation of a baseline level of assembly completeness, alongside its assumption that the LGT donor co-exists alongside the recipient in the sample under study, MetaCHIP was not effective in profiling the synthetic metagenomes introduced above. We therefore constructed a separate synthetic metagenome specifically designed for compatibility with MetaCHIP. This metagenome incorporated 20 bacterial species’ genomes each randomly spiked with 50 LGT events donated by other community members (**Methods**). These genomes were then shredded into contigs for analysis by WAAFLE and MetaCHIP, with shredded contigs derived from the same source genome provided to MetaCHIP as idealized metagenomic bins. While MetaCHIP proved highly specific when analyzing these data tailored to its approach, it detected only 17% of spiked LGT events (**Fig. S7B**). Conversely, even when conservatively penalized by a 20% database holdout, WAAFLE detected 61% of events. Hence, although WAAFLE was not specifically optimized for identifying LGT events between community members, it can do so with higher sensitivity and (unlike MetaCHIP) does not depend on binning contigs or bin completeness for LGT detection. Notably, runtimes for WAAFLE, DarkHorse, and MetaCHIP were similar with their respective upstream homology-based search steps dominating overall runtime.

### Novel LGT events across the human microbiome

To expand our understanding of LGT in human microbiomes, we applied WAAFLE to 2,376 assembled metagenomes from the expanded Human Microbiome Project (HMP1-II)^32^. After applying sample-and contig-level quality control (**Methods**), this encompassed 66 million contigs from 2,003 assembled metagenomes spanning 16 body sites and 265 individuals (**Table S1**). Among these assemblies, WAAFLE identified 116,823 contigs capturing putative LGT events (∼0.2% of all contigs above a minimum length of 500 nt). In addition to being initially identified by WAAFLE’s LGT detection algorithm, each was well-supported by read-level evidence. Specifically, each LGT junction was spanned by individual mate-pairs and/or well-covered relative to its flanking genes (**Fig. S8**). This additional evidence further helps to avoid spurious LGT calls resulting from inter-species misassembly, a hazard of real metagenomic contigs. A further 54,810 putative LGT-containing contigs (32% of initial calls) were conservatively held out of subsequent analyses due to weak read support at one or more LGT junctions (**Fig. S9**).

The final set of 116,823 LGT events detected from HMP1-II metagenomes is all, by definition, novel relative to WAAFLE’s species pangenome database. We categorized these LGT according to resolution (i.e., the taxonomic level of transferring clades), remoteness (the taxonomic level of transferring clades’ LCA), and directedness (whether or not the transferring clades could be assigned as donor and recipient, Fig. 2). Based on these definitions, 68% of putative LGT were resolved to known species, and 93% were resolved to known genera. 65% of LGT events occurred at the inter-genus level, while the remaining 35% occurred intra-genus. Only 11% of LGT calls included a clear donor and recipient clade (a value constrained by requiring longer contigs to establish directionality from gene adjacency).

**Figure 2.**
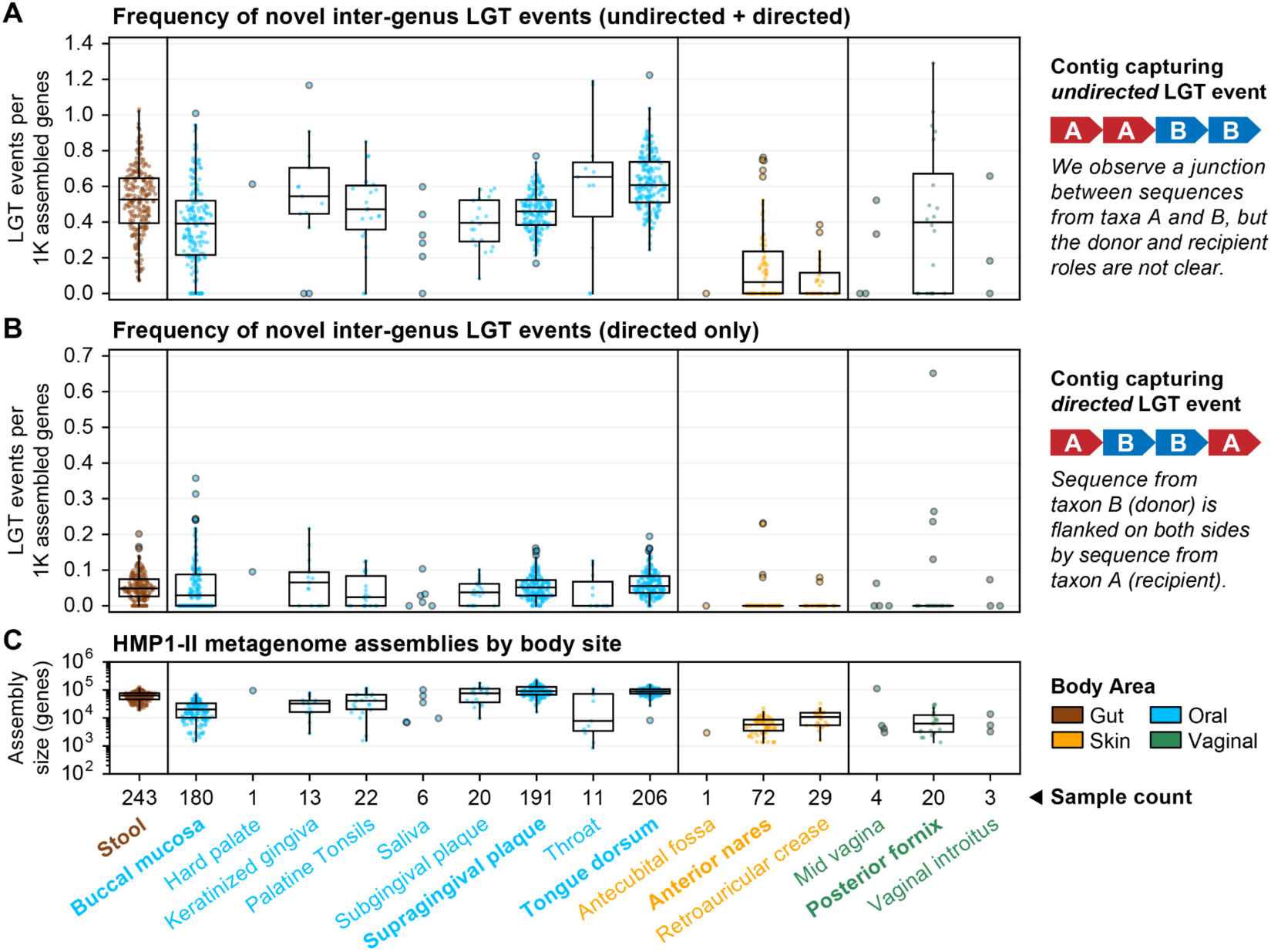
Rates of undirected and directed inter-genus LGT for HMP1-II metagenomes profiled by WAAFLE. LGT rates normalized to total assembly size in 1000s of genes, stratified by body site and correspondingly colored by body area. (**A**) All inter-genus LGT events were considered, regardless of whether the donor and recipient clades were known. (**B**) Only directed inter-genus LGT events were considered (i.e., cases where the donor and recipient clade were clear from gene adjacency). Major body sites are labeled in bold type. (**C**) Total assembly sizes for the same set of samples; only genes resolved to at least the genus level were counted. Only the first sequenced visit from each unique HMP1-II participant is plotted.

We computed overall, directed, and undirected LGT rates for whole metagenomes by normalizing total LGT events against assembly sizes (Fig. 2A). We observed that median rates of undirected inter-genus LGT were similar at the major oral, gut, and vaginal body sites: 0.4-0.6 events per thousand assembled genes. Hence, we estimate that a microbial genome from one of these environments (containing ∼2-8K genes) might be expected to show evidence for at least 1-4 novel inter-genus LGT events. We similarly computed rates of undirected LGT for clade pairs as the number of events involving the pair normalized against the pair’s total assembled gene count (detailed at species-and genus-level resolution in **Tables S2** and **S3**, respectively). Rates of directed LGT were computed similarly but normalized to the gene count of the recipient clade only (Fig. 2B and 2C, detailed in **Tables S4** and **S5**). Such rates serve as estimates of the “density” of LGT events in assembled clade pangenomes.

Rates of novel LGT between body sites’ major genera did not necessarily follow their ecological abundances (Fig. 3). For example, transfers between *Haemophilus* and *Neisseria* were consistently among the most common at oral sites, even when these genera were not among the top four by mean relative abundance. Oral genera exchanged more freely than genera from other sites, with transfers detected among all pairs of the top seven genera. Conversely, transfers in stool, anterior nares, and posterior fornix were more sparse. While sparsity at the nares and fornix sites may be influenced by their smaller assembly sizes, the same cannot be said of stool, where assemblies were comparable in size to those of oral samples (Fig. 2C). For example, while we found many LGT events between gut *Bacteroides* and *Parabacteroides* (the first- and third-most abundant gut genera), their respective transfers with *Subdoligranulum* (the fifth-most abundant gut genus) were non-existent or rare. Conversely, gut *Subdoligranulum* and *Faecalibacterium* (the sixth-most abundant gut genus) had the highest rate of LGT (∼1 event/10K genes). These findings suggest that forces beyond abundance play important roles in driving LGT rate.

**Figure 3.**
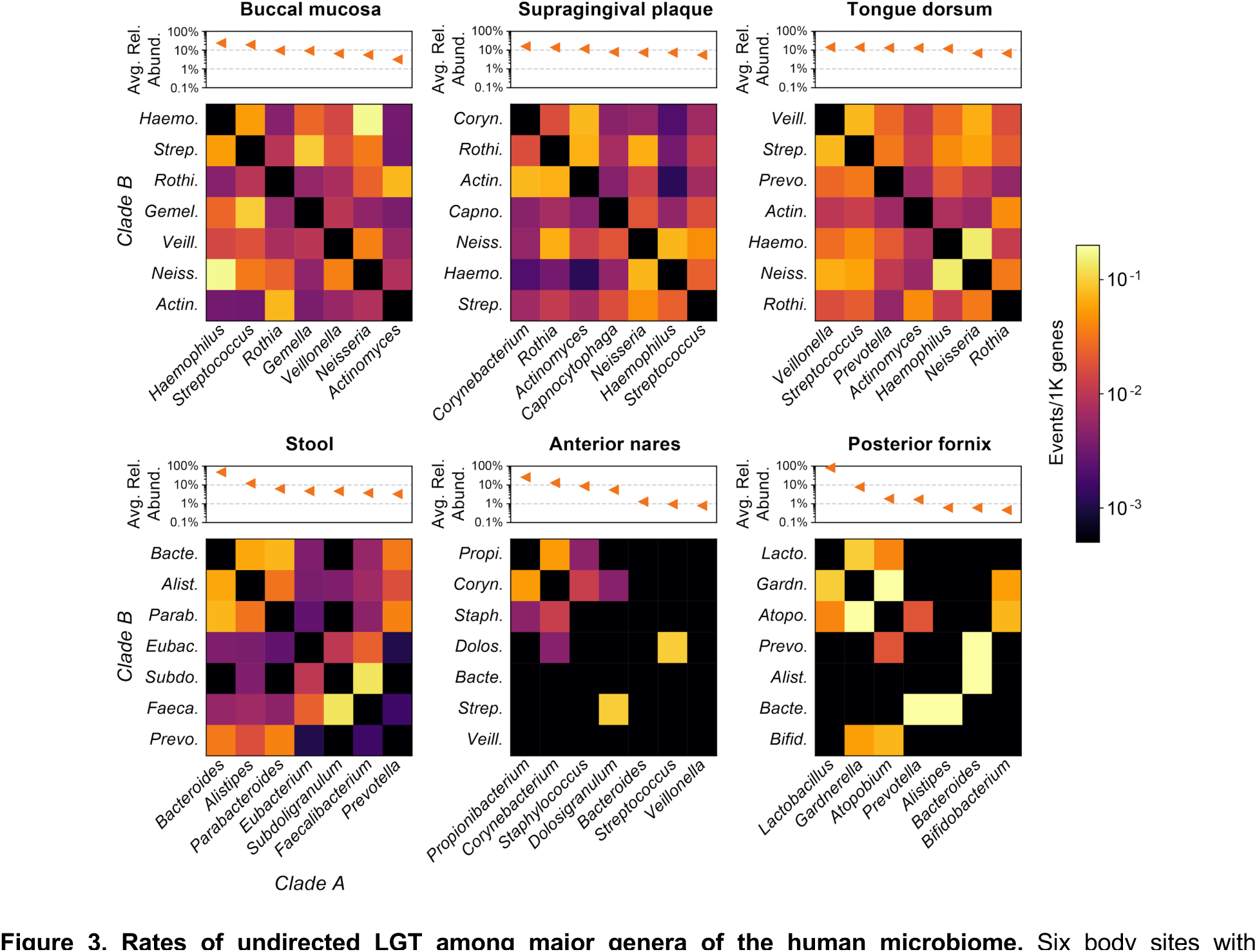
Rates of undirected LGT among major genera of the human microbiome. Six body sites with metagenome sequences from at least 20 individuals in the HMP1-II are shown. The three body sites in the top row are all from the oral cavity. Heatmap values indicate the density (rate) of undirected LGT between major genera from the body site, with “major genera” defined based on ranked average relative abundance (**Methods**). Rates are computed over first-visit samples from HMP1-II participants.

Following previously established trends for taxonomic and functional profiles of the human microbiome^32,33^, profiles of LGT presence/absence within a body site were most similar between technical replicates, followed by longitudinal samples from the same individual, then by comparisons of different individuals (**Fig. S10**). This suggests that LGT events are both unique within individuals but also prone to change over time. Notably, even profiles derived from technical replicates could be quite different, reflecting the sensitivity of assembly to the precise sampling of reads from a metagenome, particularly in the context of rare events. Indeed, this is reflected through the more comparable LGT profiles found in commonly assembled genera (those contributing 500+ genes).

### LGT rate is shaped by phylogenetic distance and donor abundance

Across six major body sites, we observed negative rank correlations between LGT rate and phylogenetic distance (PD) that were statistically significant (two-tailed *p*<0.005) outside of the nares (Fig. 4A). This finding is consistent with previous reports of enrichment for LGT between closely related species^21^ driven by similarity in genomic architecture and DNA transfer machinery^34^. Notably, our negative correlations remained statistically significant when congeneric species pairs were excluded from the analysis. This suggests that the observed trends were not solely attributable to increased LGT calls within genera and reflect a more general decreasing likelihood of LGT at increasing phylogenetic separation. Notable outliers included *Streptococcus agalactiae* with *Haemophilus haemolyticus*: a distantly diverged species pair (PD = 9.2) with a high rate of LGT inferred from plaque metagenomes (>2 events/1K genes, **Table S2**). *Streptococcus* and *Haemophilus* co-localize in the outer perimeter of oral biofilms^35^, so their physical proximity may help overcome phylogenetic barriers to LGT.

**Figure 4.**
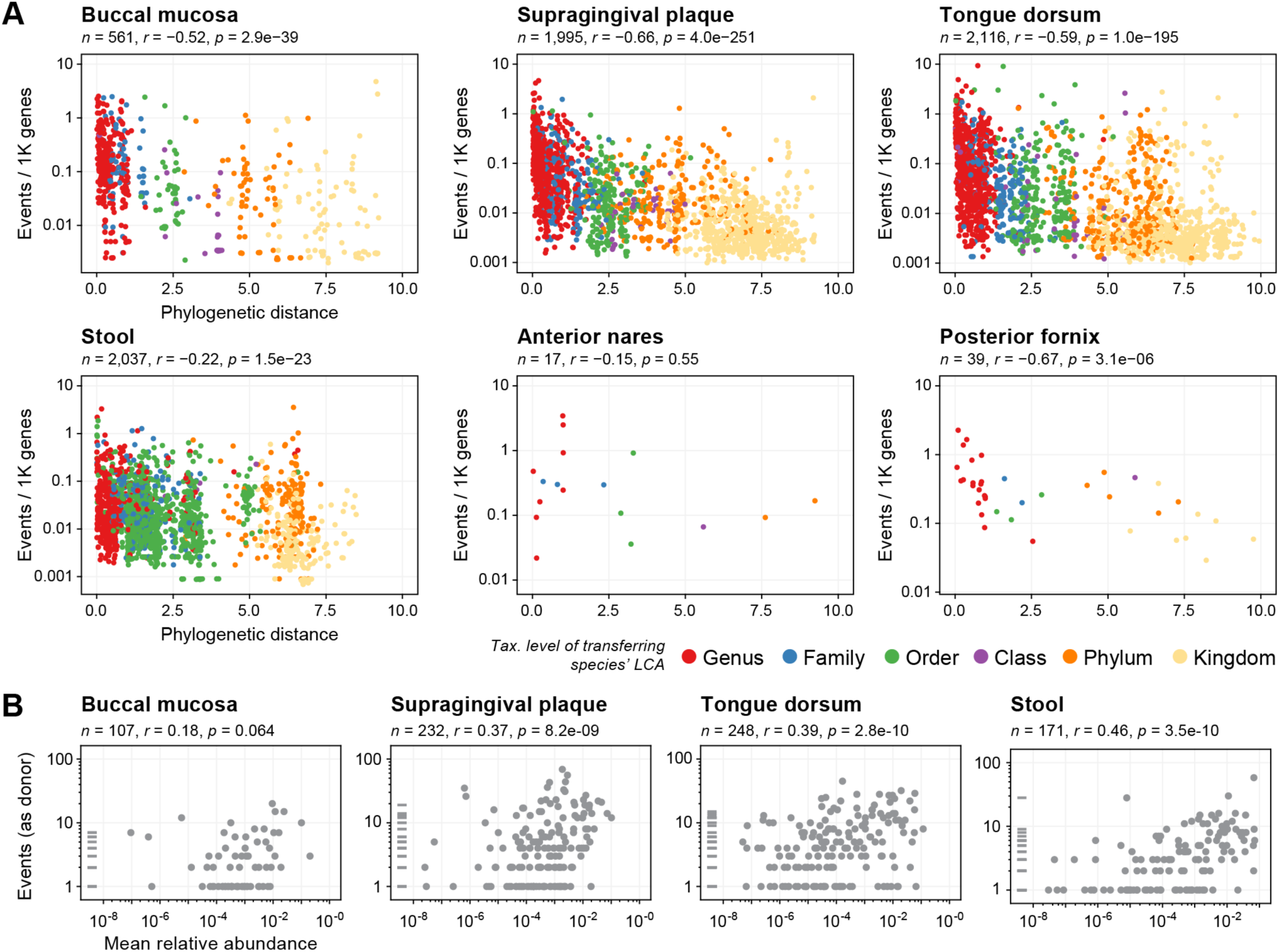
Phylogenetic distance and donor abundance as determinants of LGT rate. (**A**) Negative relationships between the density of undirected LGT for species pairs (normalized by their combined total assembly size) and phylogenetic distance at six major HMP1-II body sites. Pairs are colored according to the remoteness of the LGT (i.e., the taxonomic level of the LCA of the two transferring species). (**B**) Positive relationships between species’ frequencies as an LGT donor (inferred from directed LGT events) and species’ mean body site abundances at four major HMP1-II body sites. The nares and fornix sites were not sufficiently well represented among directed LGT and were excluded from this analysis. Horizontal marks in the y-axis margin represent species that occurred as donors, but which were never detected (i.e., having zero mean abundance). In both “(A)” and “(B)”, only species (or species pairs) contributing at least 100 genes across assembled metagenomes were considered. Correlation (“r”) values are Spearman’s rank correlation; *p*-values are two-tailed.

Phylogenetic relatedness and physical proximity both contributed to LGT rates with substantial effect, but the preceding examination of LGT among the human body’s major genera (Fig. 3) did not support a straightforward relationship between the relative abundance of a pair of clades and their LGT rate. We expanded on this by computing relationships between individual species’ abundances and their rates of LGT acquisition and donation. Critically, comparing the species abundance with the rate of LGT acquisition is complicated by the confounding effect of “assemble-ability”: that is, more abundant species tend to be better assembled, thus revealing more LGT events (**Fig. S11**). Comparing abundance with *density* of acquired LGT events (i.e., events per unit assembled genome) only partially compensated for this issue, as poorly assembled species with infrequently detected LGT exhibited a spurious negative correlation between LGT density and abundance. Trends restricted to “well-assembled” species (>10K genes per body site) were considerably flatter and never statistically significant at the *p*<0.05 level (**Fig. S12**). Hence, in this most conservative analysis, we do not find support for a relationship between species’ abundance and rate of LGT acquisition.

In contrast, we observed statistically significant positive correlations between a species’ abundance and its frequency as an LGT *donor* (Fig. 4B). Because directed LGT donated by a species are counted from other species’ genomic backgrounds, their rates of detection are not confounded with the assemble-ability of the donor species’ genomes. Examples of prolific donor taxa included *Streptococcus mitis* in the oral cavity and gut *Bacteroides vulgatus*, locally abundant species characterized by unremarkable densities of newly detected LGT events in their own genomes (**Table S2**). This difference in the effect of ecological abundance on rate of LGT acquisition vs. donation likely reflects a combination of evolutionary and biophysical forces (**Discussion**).

### Preferential attachment in the human microbiome LGT network

These trends were further evident in networks of undirected LGT events across the human microbiome (Fig. 5A). Networks exhibited clear phylogenetic organization with dense subnetworks of transfers among species within the same phylum. This was particularly evident in the gut, where transferring species pairs segregated into connected components exclusive to the Bacteroidetes and Firmicutes phyla. A similar structure was evident at the posterior fornix, with a Bacteroidetes subnetwork dominated by transfers among *Prevotella* and a Firmicutes subnetwork, including the major *Lactobacillus* species (among others). While the major genera at oral sites exchanged more freely than those of the gut (Fig. 3), species-level transfers were similarly sparse across these sites, with only 1-3% of potential species pairs from the network involved in an observed LGT event (edge).

**Figure 5.**
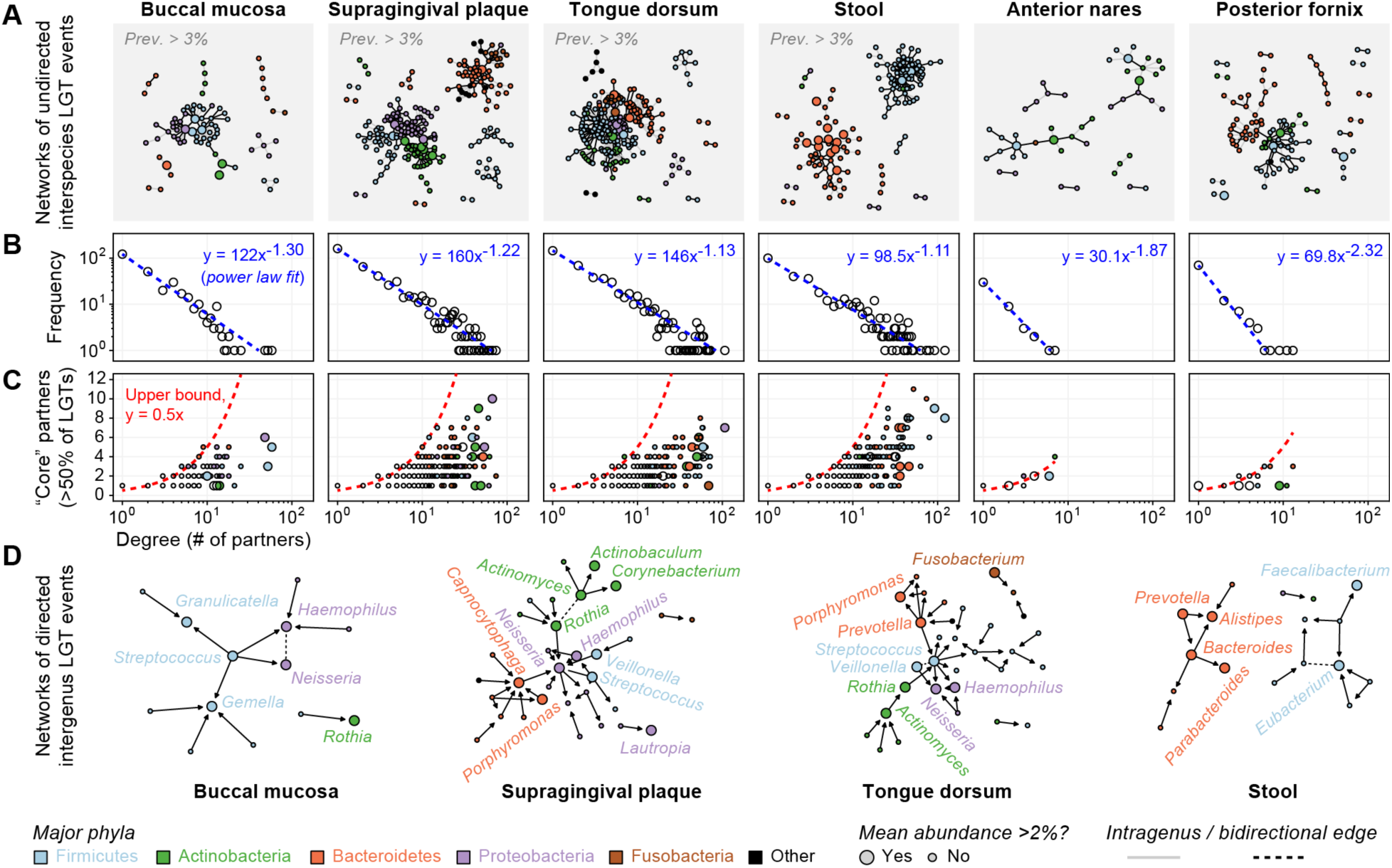
The network of LGT events in the human microbiome. (**A**) LGT shown as undirected edges between species (nodes) across six major microbiome sites. Edges from the oral and gut site are filtered for 3% population prevalence, while all edges are shown for the sparser nares and fornix sites (complete predictions in **Table S2**). Nodes are colored according to major phyla (top 5 by mean abundance) and sized according to species’ relative abundance. (**B**) Node degrees in the (unfiltered) networks from (A) follow power-law distributions, with many low-degree species and a long tail of high-degree (hub) species. (**C**) LGT events involving hub species are often dominated by a small number of LGT partners. (**D**) Directed edges are drawn from donor-to-recipient genera from the oral and gut sites. Edges are filtered for 3% population prevalence with directionality requiring at least a two-fold preference for the donor role (bidirectional edges are dashed).

This sparsity parallels an apparent “scale-free” structure of the LGT networks^36^, in which node degree (i.e. the number of LGT partners associated with each species node) followed a power-law distribution (Fig. 5B). In other words, LGT networks were characterized by large numbers of species that participated in only one or a few interactions and a smaller number of “hub” species involved in potentially many interactions with many different partners. Such patterns are common in other biological networks^37^, where they often result from preferential attachment: the tendency for a newly-formed interaction to involve a node of a higher degree. In the LGT network, this suggests that a new species joining the network is more likely to experience LGT with an existing LGT hub. This notion is consistent with our observation that abundant species tended to act as frequent donors (Fig. 4B), as we expect physical interactions with more abundant species to be more common. Conversely, species’ overall genomic plasticity did not appear to directly determine their LGT behavior, potentially because it dictates a combination of both LGT and within-species transfer events not detectable by WAAFLE.

Further, while abundant species tended to have more distinct LGT partners, they tended to favor certain ones (Fig. 5C). Indeed, the number of partners required to explain 50% of high-degree species’ LGT events was often far less than the theoretical maximum of 0.5 × degree (i.e., assuming each event occurred with a unique partner). As an extreme example, *Fusobacterium periodonticum* was involved in LGT with 69 other species across tongue samples, but 357 of 591 LGT events (60%) involved a single congeneric species, *Fusobacterium nucleatum*. This “preferred partners” phenomenon was not solely attributable to intragenus LGT and resurfaced in subnetworks containing intergenus LGT edges (**Fig. S13**). Still, the phylogenetic similarity was a comparably important factor in shaping a species’ preferred LGT partners. For example, gut *Faecalibacterium prausnitzii* was involved in transfers with 122 other species, but >50% of its LGT events involved only 8 distinct partners, all within the order *Clostridiales* (though none in genus *Faecalibacterium*).

We further analyzed the subnetwork of directed inter-genus interactions across body sites (Fig. 5D). The trends observed above were further evident in these networks, with a small number of abundant hub genera present in each. However, by layering directionality onto network edges, we were able to further characterize these hubs according to their preference for the donor or recipient role (i.e., as out-degree hubs or in-degree hubs, respectively). For example, *Streptococcus* was an out-degree hub in three oral networks, tending to donate genes at >2x the density at which it received them. Conversely, *Gemella* in the buccal site and *Alistipes* in the gut were in-degree hubs, receiving genes at greater densities than they donated. Such trends appear as asymmetries in matrices of directed LGT rates (**Fig. S14**) and were sometimes quite large, even among pairs of major genera. For example, LGT from gut *Bacteroides* to *Parabacteroides* was >10x more common than LGT from *Parabacteroides* to *Bacteroides*. Hence, in addition to preferring particular partners, clades can have a preference for the donor/recipient role in individual pairs or in the network as a whole.

### Novel LGT are enriched for mobile elements and transport functions

LGT is canonically associated with a number of molecular functions, including those that facilitate their own transfer (e.g., transposases) and those that provide a fitness advantage (e.g., antibiotic resistance). We investigated these and other functions for potential enrichment among novel inter-genus LGT. In cases where directionality was known, we examined functional enrichments in the transferred genes relative to all assembled (Fig. 6A and **Table S6**). We further examined general functional enrichments in LGT-containing contigs relative to all contigs (thus trading specificity for greater coverage, Fig. 6B and **Table S7**). Significant functional enrichments for the skin and vaginal sites were sparse due to smaller numbers of LGT assembled (Fig. 2).

**Figure 6.**
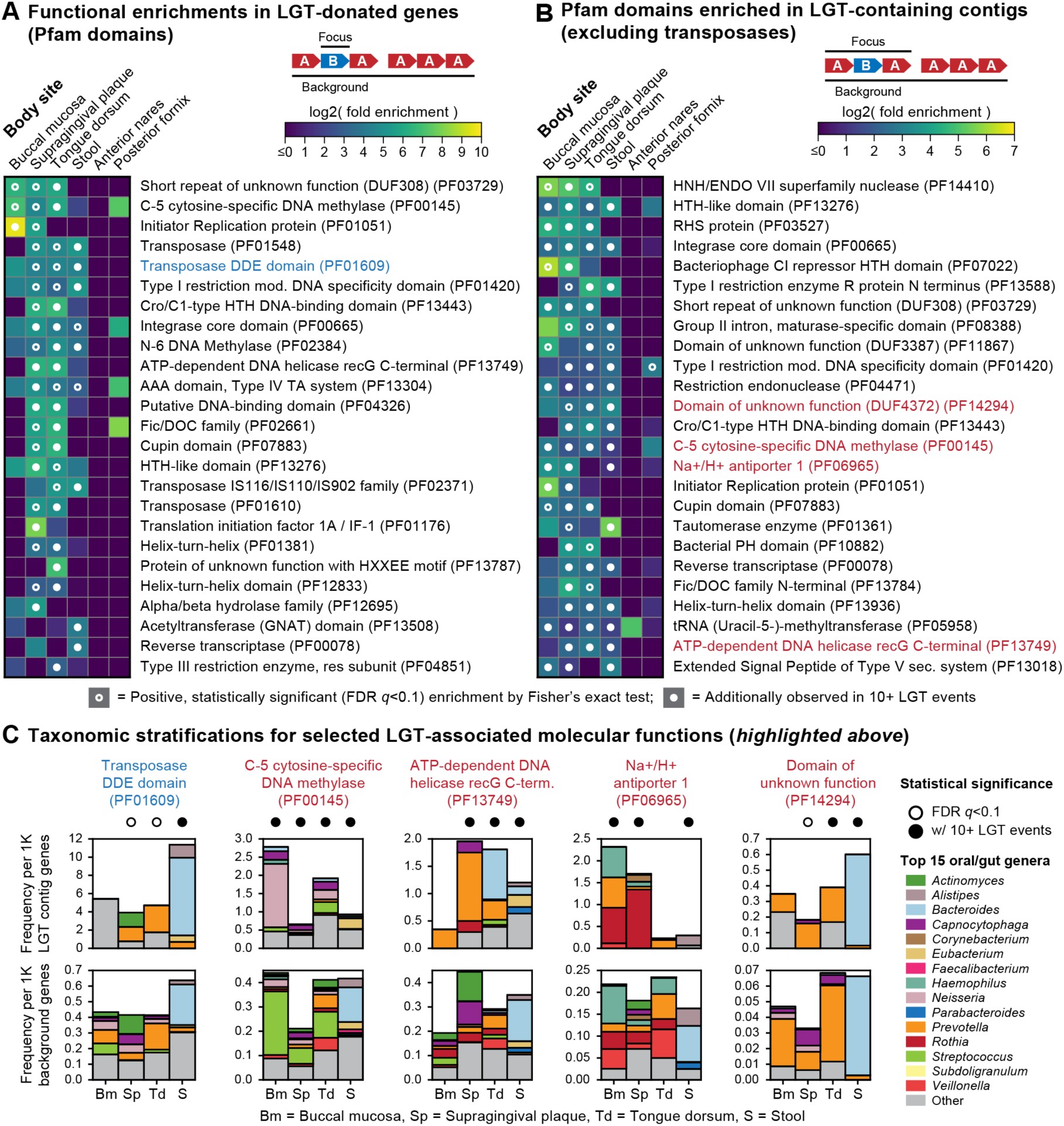
Molecular functions associated with LGT events. (**A**) Fold enrichments for Pfam^38^ domains among transferred genes from inter-genus LGT events relative to all genus-resolved genes in first visit HMP1-II metagenomes. Dots indicate statistically significant positive enrichments based on Fisher’s exact test, with nominal *p*-values subjected to FDR control (target FDR=0.1). Only domains seen in 10+ LGT events from at least one body site were considered. The top-25 such domains by mean log-scaled fold enrichment are shown. (**B**) Fold enrichments for Pfam domains among inter-genus LGT contigs relative to single-genus contigs (i.e., ignoring donor/recipient status). The top-25 domains were selected and plotted as in panel ‘(A)’, with seven transposase domains excluded to highlight other functions. (**C**) Taxonomic composition of LGT-enriched Pfam domains at oral and gut sites. The first example (blue title) is based on counts from panel ‘(A)’; all other examples (red titles) are based on counts from panel ‘(B)’.

Consistent with expectation, transposases were among the most consistently enriched functions among transferred genes (Fig. 6A). These included the Pfam^38^ transposase DDE domain PF01609 (>20x enrichment across oral and gut sites). Across body sites, transferred genes were additionally enriched for other mobile element processes, including an integrase core domain (PF00665) and a variety of more general DNA-interacting domains, including several helix-turn-helix (HTH) variants, DNA methylases, and restriction enzymes—paired components in bacterial restriction-modification systems—were also enriched. While these systems are associated with the destruction of foreign DNA (a theoretical impedance to LGT), they are selectively advantageous to their host species for defense against phage, a phenomenon proposed to have promoted their lateral dissemination^39^.

Naturally, domains enriched in LGT-containing contigs encapsulated the trends observed in transferred genes (Fig. 6B), with seven of the top-25 most enriched functions being transposases. The broader coverage of the undirected LGT dataset revealed additional trends, including enrichments for transport domains, such as Na+/H+ antiporter 1 (PF06965), and the extended signal peptide of the Type-V secretion system (PF13018), as well domains of unknown function. While the domains of unknown function could represent uncharacterized mobile elements, the transport domains do not directly interact with DNA, and hence their proliferation via LGT suggests the conferral of a fitness advantage. Antibiotic resistance was surprisingly not among the most common enriched functions, perhaps due to the stringent requirements for the HMP1-II population to be free of recent medications^32^, or alternatively, because resistance functions are already broadly represented in the pangenomes of human-associated microbes. However, we did observe a number of specific, significantly enriched antibiotic resistance domains across HMP1-II metagenomes, including *MarR* (PF01047) and *Maff2* (PF12750) at multiple oral sites and *BacA* (PF02673) in stool (**Table S7**).

Finally, we observed that the taxonomic contributions to certain preferentially transferred functions differed notably from the functions’ background taxonomic breakdown (an example of a difference in contributional diversity^40^, Fig. 6C). For example, while *Streptococcus* was a common possessor of C-5 cytosine-specific DNA methylase (PF00145) at oral sites, it was rarely involved in LGT of the function. Such genes could be involved in self-recognition by *Streptococcus* and hence of little fitness benefit in other backgrounds. Conversely, *Prevotella* were minor contributors to oral community abundance of *recG* domain PF13749 but were frequently associated with LGT of the domain. This suggests that *Prevotella*-derived *recG* improved the fitness of recipient species, leading to acquisition of the transferred gene in a variety of other genomic backgrounds.

### Experimental support for novel LGT events in human stool

We validated a subset of 21 WAAFLE-predicted LGT events from human stool using PCR amplification of genomic junctions between donor and recipient species (**Methods**). These LGT were selected from an additional dataset of 616 LGT predicted from a total of 26 assembled stool metagenomes from control participants in the HMP2 Inflammatory Bowel Disease Multi’omics Database (IBDMDB) cohort^41^, for whom biospecimens were available for experimentation. The 21 validated LGT had A+B+A+ gene adjacency, thus indicating their donor and recipient clades while also providing two junctions to validate per event, as well as short (<400 nt) donor-recipient junctions to facilitate primer design (**Table S8**).

Of the 21 LGT events under investigation, 18 received experimental support in our PCR analysis (Fig. 7 and **Fig. S15**). Of the 18 events with support, 13 showed PCR amplification of both LGT junctions (AB and BA), while 5 had weak or absent amplification at one of the two junctions, possibly due to competing sequences within the community or incompatible primer design. Validated examples include two cases of a known mobile element characterized from one clade but detected in the genomic context of another (Fig. 7A and **C**). In another example (Fig. 7B), the transferred element is of unknown function but is flanked by a pair of phage integrases, thus suggesting phage-mediated transfer. In the final example (Fig. 7D), all of the contig’s genes are uncharacterized, and the transferred gene contains a recognized domain of unknown function, which was common in LGT events (Fig. 6). Such transfers may represent uncharacterized mobile elements or potentially novel functions with adaptive significance.

**Figure 7.**
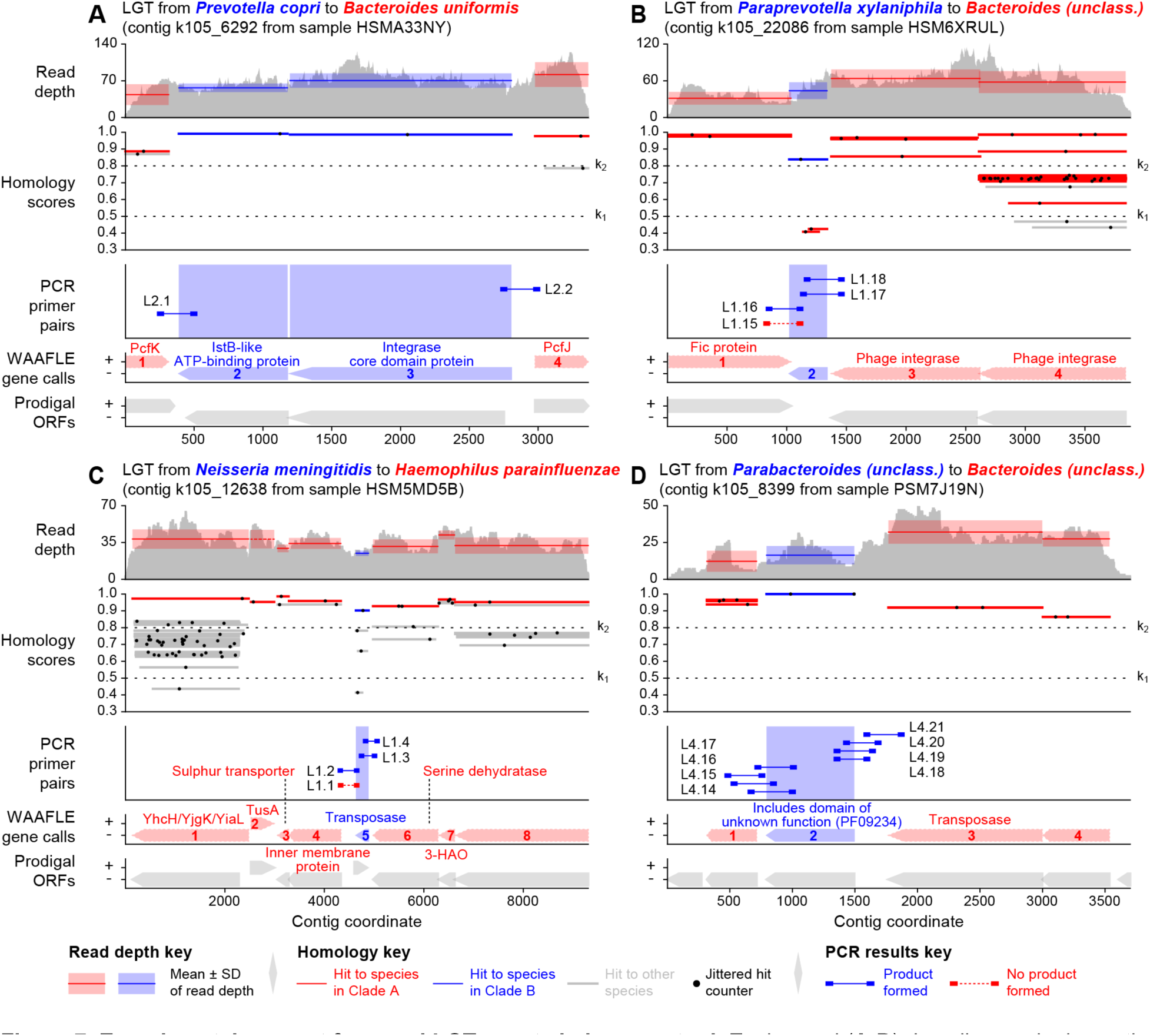
Experimental support for novel LGT events in human stool. Each panel (**A**-**D**) describes a single contig from an HMP2^41^ non-IBD control metagenome that contains a PCR-supported LGT event. “Read depth” shows variation in the depth of sample reads matching sites along the contig, as computed during WAAFLE’s quality control procedures. Bands show the mean and standard deviation of read depth for each WAAFLE-identified ORF. “Homology scores” show matches to this contig from WAAFLE’s protein-coding reference sequences, along with the k1 and k2 homology thresholds used by the algorithm (Fig. 1). Each alignment is represented by a thin gray line (indicating coverage), at a particular height (homology score), with a black dot placed randomly within the line (to facilitate counting alignments in densely aligning regions). “PCR primer pairs” shows the locations of the designed primers (line endpoints) and amplified products; some LGT events had more than one primer pair per endpoint. Lx.y refers to “Gel x, Lane y” of **Table S8**. “WAAFLE gene calls” and “Prodigal ORFs” show the location and direction of WAAFLE- and Prodigal-suggested^42^ ORFs along the contig and are largely in agreement. Features shown in red correspond to the putative recipient clade, while those shown in blue correspond to the putative donor clade. Functional annotations are taken from either 1) the UniProt-assigned^43^ name of the best homolog at each locus or 2) the UniProt-predicted domain composition of that homolog.

## Discussion

Together, these results provide substantial new insights into the landscape of LGT in the human microbiome, enabled by a novel methodology for culture-independent LGT detection and profiling in complex microbial communities. Unlike previous approaches focused on identifying LGT from sequenced isolate genomes^29,30^, WAAFLE’s focus on raw unbinned metagenomic contigs improves sensitivity and avoids the challenge of assembling complete microbial genomes from metagenomes. By applying WAAFLE to metagenomic assemblies from the human microbiome, we uncovered >100K putative LGT events across body sites, all of which were novel relative to a microbial isolate genome catalog. These findings not only highlight the vast network of species transferring genetic material within the human microbiome but also expand our understanding of LGT-based strain-level diversity and personalization. While molecular functions enriched in novel LGT events often represented mobile elements, others were of potential adaptive significance. However, in contrast with previous findings^44^, antibiotic resistance was not a dominant function among our newly detected LGT. Instead, our findings highlight the extent to which ORFs of unknown function are laterally transferred and thus likely of previously unrecognized advantage.

Expanding upon known determinants of LGT frequency, we found that not only did LGT rates between species vary inversely with phylogenetic separation, but also that spatial proximity (e.g., co-occurrence in biofilms) could overcome this. In addition, we found that a taxon’s abundance was positively correlated with its rate of LGT donation but not receipt. This trend is consistent with a physical model of LGT in which a recipient encounters the cells or free-floating DNA of abundant donors at a higher frequency. Alternatively, treating abundance as a measure of fitness, this pattern could be interpreted as an increased probability of fixation from more fit donors. These trends manifested as two distinct layers of preferential attachment: species commonly participating in LGT with abundant hub species and hubs favoring repeated LGT with phylogenetically similar partners. Surprisingly, more abundant species were not observed to *acquire* more genes via LGT. While such species have more opportunities to gain LGT, new events enter their populations at a lower frequency, which may act as an antagonistic force to fixation. These findings suggest that integrated biophysical and evolutionary modeling might be fruitful.

As a hybrid of reference- and assembly-based approaches, WAAFLE inherits their respective advantages and limitations. Notably, due to its stringent quality control filters to accommodate fragmented assemblies, WAAFLE is sensitive only to LGT events that are essentially fixed within a community. Loci containing potential LGT with coverage differing from adjacent regions—which could occur during a sweep prior to fixation—cannot be reliably distinguished from assembly errors and are excluded, potentially rejecting LGT events that are specific to one lineage (e.g., a strain) or not fixed in the population. In addition, while WAAFLE does not directly map reads to a reference database, it is still dependent on a reference catalog of microbial pangenomes for taxonomic and functional annotation of metagenomic contigs. Incompleteness within this catalog can potentially lead to spurious LGT calls: for example, if a gene is core to a given genus but deleted from the single reference genome of a species X within that genus, WAAFLE may consider the gene to have been acquired by interspecies LGT when observed in new metagenomic strains of X (WAAFLE’s ambiguity and sister-genome filters conservatively remove such cases where possible). Conversely, because WAAFLE is focused on detecting new instances of genetic material entering a species’ pangenome, by design it will not highlight known (and potentially ancient) LGT events within the pangenome when they are re-detected in novel metagenomic strains of the species. This limitation could be addressed in future versions of the software by adding known LGT events as a new layer of pangenome annotation. More generally, future analyses with WAAFLE will benefit from improved reference genome catalogs, specifically from recent efforts to expand the catalog of human-associated microbial genomes^45–50^.

Because WAAFLE uses metagenomic contigs as input, it is not sensitive to the general challenge of assembling complete genomes from metagenomes. That said, future versions could also be re-tuned for application to metagenomic assemblies of increasing quality or to utilize explicit pre-defined taxonomic binnings, which may aid in disambiguating recipient and donor taxa among candidate LGT embedded within short contigs. WAAFLE is ultimately most sensitive to the reliability of individual contigs and will benefit from new methods to limit or identify misassembly events, including improvement in metagenomic assemblers themselves^51^, as well as downstream filtering methods expanding the junction-coverage approach implemented here^15^. In the future, these limitations of assembly could similarly be bypassed entirely by applying WAAFLE to long-read sequencing data^52^, as long (multi-kilobase) reads share the multi-gene “scope” of short metagenomic contigs still fall short of lengths required by existing LGT methods, and may avoid misassembly issues.

While we acknowledge these limitations inherent to our current implementation of a contig-based LGT profiler, we note that our attempted benchmarking efforts of other potentially suitable classical sequence-based LGT detection methods were either not successful (e.g., Alien_Hunter) or highlighted comparatively poorer LGT detection (e.g., DarkHorse). Additionally, we showed that WAAFLE outperformed MetaCHIP^31^, an explicit phylogenetic LGT profiler for detecting within-community LGT, using synthetic benchmarking data. Still, WAAFLE does not incorporate a phylogenetic validation of candidate LGT, suggesting an ongoing and complementary role for methods like MetaCHIP. While many trends were evident from our initial analyses of novel LGT in the human microbiome, much remains to be uncovered. Further investigation of the mechanisms of transfer and fixation of LGT-enriched functions is warranted, particularly those associated with uncharacterized domains or not obviously attributable to mobile element processes. Finally, while WAAFLE-identified LGT are, by definition, novel, more work is needed to formally establish their age. While some candidate events are likely ancient, low levels of adaptation to the recipient genomes, coupled with evidence of stable personalization of LGT across participants here, suggest that novel LGT events arise and fix within individual human microbiomes as part of their long-term developmental dynamics^3,53^, potentially influencing host health and disease.

## Methods

We analyzed and experimentally validated putative LGT events from the HMP1-II^32^ and HMP2^41^ populations, predicted by the herein newly developed method (WAAFLE) to identify novel LGT events from assembled metagenomes. This section describes the HMP1-II and HMP2 metagenomes, including computational and experimental analyses of newly predicted LGT events, as well as the WAAFLE algorithm, implementation, its validation, and comparisons between WAAFLE and other LGT-detection methods. Additional details can be found in the Supporting Information. Input data, software implementations, tutorials and vignettes, and analysis results are available at http://huttenhower.sph.harvard.edu/waafle.

### The WAAFLE algorithm

The WAAFLE algorithm consists of two steps: 1) a homology-based search of the input contigs against a gene sequence database followed by 2) taxonomic classification of contigs, which includes LGT identification. An optional intermediary step (1.5) identifies candidate protein-coding loci (potential ORFs) along the input contigs from the results of the homology-based search. An optional final step (3) maps sequencing reads back to taxonomically annotated contigs to aid in the identification and exclusion of misassembly events.

#### Step 1: Homology-based search

In its current implementation, WAAFLE uses BLAST^54^ to perform nucleotide-level search of input contigs against a gene-sequence database. The sensitivity of *blastn* was retained in favor of accelerated options since a wide range of homology detection is useful, and the relatively small size of assembled metagenomes prevents this step from being a computational bottleneck. WAAFLE’s *blastn* call uses default search options and requests the fields from tabular output format “6” with the following exceptions: 1) WAAFLE raises the default number of database hits per query to 10K (required for improved taxonomic coverage for larger contigs); and 2) WAAFLE additionally requests the inclusion of query (contig) length, subject (database sequence) length, and subject strand in the output fields. This step is governed by the *waafle_search* script within the WAAFLE package.

WAAFLE scores alignments (“hits”) of a contig against a gene sequence in its database according to the product of 1) alignment percent identity and 2) modified subject coverage (mSC). We refer to this product as a WAAFLE “homology score.” Like traditional subject coverage, mSC captures the fraction of the gene (database) sequence that was covered by a contig in its local alignment. However, unlike traditional subject coverage, mSC does not penalize gene length that falls beyond the endpoints of the contig, as would occur in the case of a partially assembled gene. For an alignment to the positive strand of the database sequence, mSC is defined as:

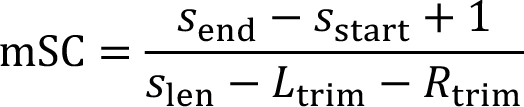

Where *s*_end_ and *s*_start_ are the start and stop coordinates of the alignment within the gene sequence, *s*_len_ is the gene’s length, and *L*_trim_ and *R*_trim_ are adjustments for alignments involving the left and right end of a contig (respectively), defined as:

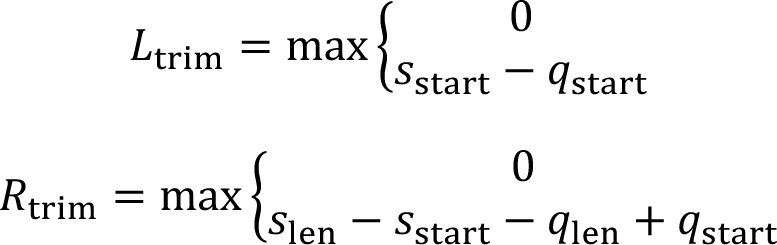

Where *q*_start_ and *q*_len_ are the start coordinate of the alignment within the contig and contig length, respectively. Note that when *L*_trim_ and *R*_trim_ are both 0, the contig could (in principle) contain an alignment to the full length of the gene, and so mSC reduces to traditional subject coverage. For alignments to the negative strand of the database sequence, mSC is calculated as above, with *s*_start_ replaced by *s*_len_ - *s*_start_ + 1 and *s*_end_ replaced by *s*_len_ - *s*_end_ + 1 (i.e. their corresponding 1-based positions on the positive strand of the database sequence).

#### Step 1.5: Identifying candidate protein-coding loci

This optional step can approximate the coordinates of protein-coding ORFs (loci) within the input contigs (as an input to Step 2) based on the results of the homology-based search. This step is governed by the *waafle_genecaller* script. Alternatively, users can provide independently called ORFs in GFF format for Step 2 (below). Any analyses in this study that did not use WAAFLE’s intrinsic gene calling used Prodigal^42^ for this purpose.

To identify candidate protein-coding loci along a contig from hits to the gene sequence database, hits with poor mSC are first excluded (tunable default, mSC<0.75). Then, a network is constructed from all hits to the contig with edges connecting hits that overlap above a target threshold. The overlap is defined as the fraction of the shorter hit contained within the longer hit, and the threshold is conservatively set to 0.1 (tunable parameter). WAAFLE then isolates the connected components of this network (i.e. collections of overlapping hits) as candidate loci. The rightmost and leftmost endpoints of hits within the connected component are taken as the start and end coordinates of the corresponding coding locus. Short loci are then filtered (tunable default, loci <200 nt in length removed), and the remaining loci are written to a GFF file. This step can optionally be run in a strand-aware mode that will only connect hits on the same strand of the contig (positive or negative). By default, the strand of the longest hit contributing to a candidate locus is taken as the strand of the candidate locus.

We emphasize that this aspect of the WAAFLE algorithm is provided as a convenience. WAAFLE’s candidate protein-coding loci are not intended or expected to outperform those of a dedicated ORF-calling program, such as Prodigal. That said, in practice, WAAFLE locus calls and Prodigal ORF calls tended to agree well (Fig. 6 and **Fig. S16**).

#### Step 2: Taxonomic classification of contigs and candidate LGT identification

In Step 2, results of the homology-based search are combined with the coordinates of putative protein-coding loci (“loci”) within contigs to 1) taxonomically classify the contigs and 2) identify putative LGT-containing contigs. This step of the WAAFLE algorithm is conducted separately for each contig and involves a number of subroutines. The subroutines act on 1) the contig; 2) the collection of hits to the contig, with taxonomic and functional annotations inferred from the subject sequence name; and 3) the taxonomy file relating species included in the gene sequence database. This step is governed by the *waafle_orgscorer* script within the WAAFLE implementation.

##### Subroutine 2a: Locus scoring

A homology score is computed at each locus *l* in contig *L* for each species *s* in the total set *S* whose genes are aligned to the contig. When a hit to a gene from species *s* overlaps with locus *l* covering one or more nucleotide positions, the homology score of the hit is assigned to each position. If a subsequent hit from species *s* covers an already-hit position in *l*, the better of the two scores is saved. After all hits are processed, the homology score for species *s* at locus *l* is the average of the scores assigned to each nucleotide position, treating unhit positions as having a value of 0. After this initial scoring, loci that never exceeded a target homology threshold for any species (pre-optimized default, k_1_ = 0.5) are masked and ignored. WAAFLE implements additional parameters to filter low-quality hits and genes at this stage, if desired. The end product of this stage of the algorithm is an |*S*|×|*L’*| matrix *H* of per-species, per-locus homology scores, where *L’* is the set of unmasked loci.

If using a functionally annotated gene sequence database, functions are additionally transferred to loci at this stage based on the results of the homology-based search. Specifically, for each functional annotation system (by default, UniRef90 and UniRef50^55^), the annotation(s) of the best-scoring database sequence are transferred to their hit loci, independent of sequence taxonomy. These transfers can be additionally filtered by requiring the best hit to exceed a given homology threshold (the predefined threshold k_1_ = 0.5 is used by default).

##### Subroutine 2b: Evaluating single-clade explanations

WAAFLE first attempts to find a single species A to which the contig can be attributed. To “explain” the contig, the species must have a homology score for each locus along the contig that exceeds a given lenient threshold, k_1_ (with a pre-optimized default of 0.5, Fig. 1A). Symbolically, given the |*S*|×|*L’*| scoring matrix *H* (defined above), a species A explains the contig if:

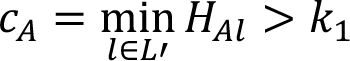

Where *c*_A_ is the one-clade “critical” score for species A. If multiple species have critical scores exceeding k_1_, then each species is ranked by its average per-locus score, defined as *r*_A_ for species A as:

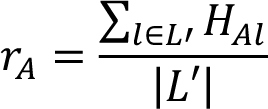

Any species within “range” 0.05 (tunable parameter) of the best score are melded, and the contig is assigned to the LCA of those species. If one species has a singularly best score, the contig is assigned to that species. In either case, the algorithm halts.

##### Subroutine 2c: Evaluating two-clade (LGT) explanations

If a single-species explanation for the contig cannot be found (i.e., *c*_s_ < k_1_ for all *s* in *S*), WAAFLE then tries to find an explanation for the contig involving LGT. An LGT-based explanation requires that a pair of species (A, B) confidently explains each locus along the contig, meaning that at least one of the two species has strong homology to each locus (i.e., a homology score exceeding a stringent threshold, k_2_, with a pre-optimized default value of 0.8; Fig. 1A). Symbolically, we require:

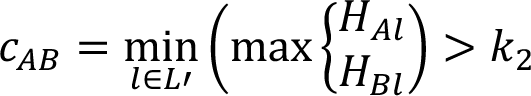

Where *c*_AB_ is the two-clade critical score for the species pair (A, B). Each species pair is additionally ranked by a similarly defined average score (*r*_AB_):

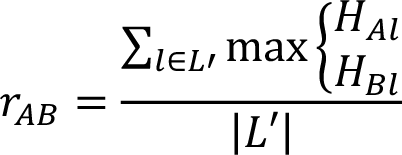

If one or more species pairs’ critical scores exceed k_2_, then the contig is considered a candidate LGT. Among such pairs, any with an average score within range 0.05 of the best average score are saved as “equivalently good” explanations for the LGT.

##### *Post hoc* filtering of candidate LGT

WAAFLE applies *post hoc* filters to candidate LGT to rule out alternative explanations, e.g., gene deletion in the putative recipient. First, a candidate LGT between species A and B is rejected if too large a fraction of the contig could be assigned to either species A or B (pre-optimized default, 10% of sites). The candidate LGT is additionally rejected if a gene *g* that was putatively transferred from B to A was also observed in a sister clade of A (A’). Such evidence supports an alternative explanation for the contig in which gene *g* was present in the parent of A and A’ and deleted along A’s lineage.

##### Assigning LGT directionality

If a single species pair (A, B) explained the contig and passed both of these filters, then the contig is reported as an LGT event between A and B and the algorithm halts. If loci assigned to species A flank both sides of the loci assigned to species B (i.e., an A+B+A+ adjacency pattern in regular expression syntax), then the LGT is considered a directed LGT from B (the donor) to A (the recipient). If multiple species pairs explained the contig and passed the above filters, WAAFLE attempts to meld these pairs into a single explanation for the contig (following the “melding” procedures introduced in Subroutine 2b). If the pairs require different gene adjacency configurations to explain the contig (e.g., AABA and ABAA), the melding process fails and WAAFLE rejects the candidate LGT and proceeds to the next stage of the algorithm. Otherwise, the donor and recipient species are individually melded to their respective LCAs. If one of the resulting LCAs is a descendent of the other, the melded explanation is also rejected.

##### Taxonomic iteration

If WAAFLE fails to find a single-species or species-pair (LGT-based) explanation for a contig, it repeats subroutines 2a, 2b, and 2c at the genus level (Fig. 1A). In subroutine 2a, homology scores are assigned between contig-loci and genera by regrouping species-level hits according to their parent genera in the input taxonomy. This iterative process is repeated until 1) an acceptable one- or two-clade explanation is found for the contig or 2) the root of the taxonomy is reached. In the latter scenario, the algorithm halts and the contig is reported as “unclassified.” Users can optionally perform one or more “jumps” up the taxonomy before attempting classification (e.g., to ignore intra-genus LGT events).

#### Step 3: Coverage-based quality control

A final optional but recommended step in the WAAFLE workflow performs further quality control and filtering of candidate LGT by ensuring even coverage and reliable junctions at putative LGT sites. This includes mapping all sequencing reads (including unassembled) to their corresponding metagenomic contigs (**Fig. S8**). This step is governed by the *waafle_junctions* and *waafle_qc* scripts within the WAAFLE package.

Mappings of sequencing reads to LGT-containing contigs are used to support or refute candidate LGT in two ways. First, WAAFLE looks for sequencing fragments that span the gene-gene junctions supporting an *LGT junction* (i.e., the transition or space between a gene from the donor species to the recipient species). Such fragments provide strong support that the junction is a biological structure and not the result of misassembly. However, for junctions that are larger than the size of typical sequencing fragments (e.g., a few hundred nucleotides for typical Illumina HiSeq libraries), it will not be possible to achieve this type of LGT support.

As a second check, the average coverage of the LGT junction is thus compared to the average coverage of the two flanking genes from the donor and recipient species. If coverage of the LGT junction is at least half that of the flanking genes, WAAFLE considers this as reasonable support for the LGT. If at least one of the junctions flanking an LGT event fails both of these checks (i.e., no sequencing fragments span the junction and the coverage of the junction is weak relative to the coverage of the flanking genes), then the LGT is considered dubious.

### WAAFLE validation and evaluation

We optimized and benchmarked WAAFLE using synthetic contigs containing LGT at different levels of taxonomic remoteness as well as control contigs with no LGT. True positive rate (TPR) was calculated as the fraction of LGT-containing contigs identified to contain a LGT event by WAAFLE (separately for LGT of increasing taxonomic remoteness, Fig. 1). False positive rate (FPR) was calculated as the fraction of control contigs erroneously identified as containing an LGT. The stringency of the FPR measure was scaled based on the allowed minimum taxonomic remoteness of the transferring clades. For example, the most stringent FPR measure considers any interspecies LGT call for a control contig to be a false positive, while a more relaxed LGT measure ignores false positive calls at the intragenus level, the intrafamily level, and so forth (Fig. 1). We further evaluated the accuracy of the taxonomic assignments made for correct LGT and control (no LGT) calls. This evaluation was stratified according to recipient/control taxon and donor taxon and quantified the fraction of correct calls at each taxonomic level (kingdom, phylum, class, order, family, genus, and species, **Fig. S6**). The evaluation procedures were additionally repeated while holding out fractions of WAAFLE’s sequence database (5, 10, and 20% of genes within each pangenome) to simulate uncharacterized species pangenome diversity.

#### WAAFLE reference data

WAAFLE uses as a reference a collection of protein-coding gene sequences and a species taxonomy; these are provided with its implementation. This is currently the ChocoPhlAn gene family database produced during the development of MetaPhlAn2^56^ and bundled with HUMAnN2^40^. The individual coding sequences in this database were annotated against UniRef90 and UniRef50^57^ by homology-based search during HUMAnN2 development. Here, we used UniRef annotations to link out to other functional annotation systems (e.g. Pfam^38^ domains) via UniProt^43^. A *blastn*-formatted version of this sequence database and corresponding taxonomy file are available from http://huttenhower.sph.harvard.edu/waafle. Additional format details of the sequence database and taxonomy file are available from the WAAFLE user manual, which is linked to the preceding URL.

#### Construction of synthetic contigs

We constructed synthetic contigs by concatenating protein-coding sequences (“genes”) from reference genomes included in WAAFLE’s underlying pangenome database. Each synthetic contig dataset was based on a target adjacency pattern (*p*), a fixed number of species pairs (*n*), a fixed number of LGT events per species pair (*m*), and a decision to optionally end-truncate the contig after gene concatenation. The adjacency pattern *p* is a string of the characters “A” and “B” representing the ordering of source genes from the recipient (A) and donor (B) species, respectively. For example, the pattern “ABA” represents a single species B gene flanked by two species A genes (representing a B-to-A transfer). For each level *t* of the taxonomy, we randomly selected *n* pairs of species (A, B) whose taxonomic LCA occurred at level *t*. For example, at the family level, the species *Bacteroides fragilis* and *Acetothermus paucivorans* would be a candidate LGT pair as their taxonomies diverge at the family-level clade Bacteroidaceae.

For each of the *n* species pairs (A, B), we randomly selected one species A reference genome and one species B reference genome from which to sample genes for contig generation. To construct a single contig for the species pair (A, B) with adjacency pattern ABA, we would first randomly select a gene from the A reference genome, then concatenate this gene with a randomly selected gene from the B genome, followed by a final randomly selected gene from the A genome. Gene sequences selected for concatenation were randomly mutated at 5% of nucleotide sites to simulate homologs from new isolates of species A and B. We excluded genes <200 nt in length during random sampling. To simulate partially assembled genes, some contigs were randomly truncated within their 5’ and 3’ genes (but always leaving at least 200 nt of truncated gene within the contig). For a given adjacency pattern and level of taxonomic remoteness, we repeated this step *m*=40 times for each of *n*=250 species pairs to produce a synthetic dataset of 10K LGT-positive contigs. We produced similar datasets for a variety of adjacency patterns (Fig. 1) with taxonomic remoteness at the kingdom, phylum, class, order, family, and genus levels.

We constructed control (negative) contigs following similar procedures. However, instead of sampling genes from pairs of species, we sampled genes from pairs of reference genomes (A, B) belonging to the same species. Thus, the control contigs are identically constructed to our synthetic LGT contigs except for the fact that all of their genes derive from the same species pangenome rather than two distinct species. Positive and negative control contigs are available for download from the WAAFLE website (http://huttenhower.sph.harvard.edu/waafle).

#### Optimizing gene-calling parameters

WAAFLE’s method for finding candidate protein-coding loci (Step 1.5) in a contig based on hits discovered during homology-based search (Step 1) involves three tunable parameters: 1) the minimum mSC for a hit to be used in locus definition, 2) the minimum overlap for grouping hits as part of a connected component in the developing locus graph, and 3) the minimum size for a reported locus. We evaluated a range of settings for these parameters by applying WAAFLE’s gene-finding procedure to synthetic contigs containing partial genes at their 5’ and 3’ ends (“end-truncated contigs”, **Fig. S16**). We considered a gene call to be a true positive (TP) if it uniquely covered a known gene on the synthetic contig with >80% mutual coverage and a false positive (FP) otherwise. Known genes that were never covered in this way were counted as false negatives (FNs). For these evaluations, we computed the true positive rate [TPR = TP/(TP+FN)] as a measure of sensitivity and positive predictive value [PPV = TP/(TP+FP)] as a measure of specificity. These measures avoid the need to specify “true negative” (TN) gene-coding loci, which are challenging to define (e.g., an intergenic space could be interpreted as a single TN or multiple TNs).

WAAFLE performed well across parameter settings, with TPR values ranging from 0.64 to 0.96 and PPV values ranging from 0.95 to 0.99. Loss of sensitivity was primarily explained by the exclusion of shorter genes (<200 nt in length) or similarly sized gene fragments at the ends of contigs. When tuned to allow shorter genes in its output, WAAFLE performed similarly to Prodigal^42^ (a dedicated ORF caller, **Fig. S16**). However, because very short genes were ultimately problematic for LGT detection, by default, WAAFLE’s gene caller imposes a minimum 200-nt filter on candidate gene-coding loci during the optional gene-calling step. (As discussed in the next section, similarly short genes are ultimately filtered prior to LGT calling even if a user provides their own independently generated ORF calls.) While varying the overlap parameter had only a small effect on accuracy, higher values of the minimum mSC parameter (including WAAFLE’s conservative default of 0.75) traded specificity for sensitivity.

#### Optimizing LGT-calling parameters

We selected and optimized tunable parameters for WAAFLE by initially assigning each parameter to an intermediate default value within its potential range. We then optimized parameters one-at-a-time by analyzing sets of synthetic contigs using different settings of the parameter within its allowable range. We focused on two synthetic contig configurations for this work: AB-adjacency contigs that were truncated at both ends, the minimal case of LGT that WAAFLE could theoretically detect, and AAAAABAAAAA contigs, which reflect a more idealized LGT-containing contig from a partially assembled metagenome. We refer to these as “short” and “long” contigs for simplicity below. Benchmarking was conducted with 20% of species pangenomes held out during homology-based search: a harder, but more biologically realistic, scenario simulating incomplete understanding of species’ pangenome content.

##### Gene filtering parameters

Some parameters were more impactful for detecting LGT in short contigs and others in long contigs. WAAFLE’s defaults attempt to balance performance across these scenarios. Parameters associated with handling weakly supported candidate gene loci (“loci”) impacted performance on both contig types. For example, the option to assign an “unknown taxon” when a given locus lacked reasonable homologs in any species resulted in a large spike in false positive rate (FPR, **Fig. S5**). However, penalizing such loci (by still forcing them to be “explained” by a known species) dramatically reduced the true positive rate in longer contigs. Surprisingly, simply ignoring such loci (i.e., treating them as probable misannotations and not requiring species to explain them) provided the best balance of sensitivity and specificity. In addition, both long and short contigs benefitted from a filter on minimal gene length, as short (likely miscalled) loci with random homology to a known gene would be erroneously identified as LGT events. The default threshold of 200 nt covers 94% of previously defined genes in WAAFLE’s underlying reference database.

##### Homology threshold parameters

WAAFLE’s k_1_ and k_2_ parameters (introduced above) define the minimum homology scores that a single clade or pair of clades must achieve over a contig’s gene-coding loci to assign the contig to that clade (or pair of clades). WAAFLE’s default homology score, percent nucleotide identity × modified subject coverage (mSC, defined above), falls in the interval [0, 1]. An average score of “1” over the length of a candidate gene-coding locus from an input contig indicates that the (potentially partially assembled) locus exactly matched a reference gene over its entire length.

The value of k_1_ had little influence on the accuracy of LGT calls for short contigs, while the value of k_2_ had a small influence on FPR in small contigs (with larger values reducing FPR, **Fig. S2**). Thus, the tuning of k_1_ and k_2_ was largely driven by performance on longer contigs, where they induced greater variation in accuracy. Larger values of k_1_ made it harder to assign a longer contig to a single clade. This had the effect of improving sensitivity in cases where the LGT-recipient contained a remote homolog of a transferred gene (the remote homolog fails to score well enough to participate in a single-clade explanation for the contig, allowing the LGT-based explanation to be considered). However, the same principle caused a larger fraction of correct single-clade explanations to be ignored in favor of LGT-based explanations, resulting in elevated FPR. Hence, a conservative default of k_1_=0.5 was selected (erring on the side of missing true LGT in favor of one-clade explanations).

Conversely, higher values of k_2_ made two-clade (LGT) explanations harder to accept, resulting in decreased TPR and FPR. Thus, the default setting of k_2_=0.8 was also selected to be conservative while avoiding a marked loss of TPR for inter-genus LGT events that occurred at k_2_>0.8. Like all parameters in WAAFLE, k_1,_ and k_2_ can be fine-tuned by the user to favor sensitivity (by increasing k_1_ and/or decreasing k_2_) or specificity (by decreasing k_1_ and/or increasing k_2_).

##### Parameters for filtering candidate LGT

The majority of WAAFLE’s remaining parameters are responsible for post-hoc filtering of candidate LGT that passed the two-clade homology threshold. As introduced above, these include options for removing LGT that could be alternatively explained by gene deletion, as well as options for melding solutions involving multiple pairs of clades.

LGTs are initially filtered on the fraction of “ambiguous” sequence along the length of the contig (i.e., explainable by both the donor and recipient clades). The allowed fraction of ambiguous sites is tunable, with larger fractions tending to allow more false positive events, mostly at the intra-genus level (the default is set conservatively to 10%, favoring specificity, **Fig. S3**). The threshold for calling a gene “ambiguously explained” can be set to 0 (off), k_1_ (lenient), or k_2_ (strict), which all produced similar results. The “sister-clade filter” introduced above (as part of Subroutine 2c) was additionally found to reduce false positive intra-genus calls at almost no cost to sensitivity, even at its stricter setting (i.e., treating even remote homologs of putatively transferred genes in sisters of the putative recipient clade as disqualifying, **Fig. S4**).

Finally, we tuned several parameters related to the “melding” of alternative LGT-based explanations for a contig involving multiple species pairs. Curiously, we found that reporting the best-scoring solution when multiple solutions were found was not optimal, as the presence of multiple explanations was itself associated with false positive LGT calls (**Fig. S4**). This was particularly true for explanations involving e.g., a clade A and either clades B or B’, where the LCA of B and B’ was also the LCA of A, B, and B’. Combinations of parameters that find and reject such solutions improved specificity (albeit slightly) with no cost to sensitivity.

### Comparison with other methods

Prior methods and tools for LGT detection have mostly been designed to find LGT events in sequenced isolate genomes^58^ by identifying islands of unusual sequence composition compared to background^29^ or by finding discordance between species trees and gene trees^59^. Regrettably, such methods are prone to underestimating LGT diversity in microbial communities. We set out to compare WAAFLE with previously published tools for LGT identification: alien_hunter^26^, DarkHorse^27^, and MetaCHIP^31^. Initial evaluations with alien_hunter indicated that it would not be appropriate for metagenomic contigs due to the algorithm’s minimum input-sequence length requirement (a conclusion later confirmed in writing by the method’s developers). DarkHorse, on the other hand, was amenable to application and evaluation involving the same sets of synthetic contigs used for WAAFLE benchmarking, while a robust comparison to MetaCHIP would eventually require the construction of a new, more compatible synthetic dataset (described later in this section).

#### WAAFLE vs. DarkHorse

DarkHorse uses a reference-based homology search and integrates BLAST scores and phylogenetic statistics of a genome to produce global probabilities of the entire genome where sequences (genes) with lower probability scores are candidate LGT events. We ran DarkHorse (v2.0) following recommendations from its online manual (http://darkhorse.ucsd.edu/README.txt). Specifically, we first used Prodigal^42^ (v2.6.3) in “`-p meta`” mode to identify and translate ORFs along the synthetic contigs. We then executed DarkHorse’s initial search (the equivalent of Step 1 from the WAAFLE algorithm) by aligning the translated ORFs to DarkHorse’s database. This search used DIAMOND^60^ (v0.9.19.120) as a protein-level alignment engine with the suggested flags “*`-e 1e-3 and --max-target-seqs 100`*.” We then divided the alignment results according to query contig and performed the second step of DarkHorse analysis -- i.e. classification with the `darkhorse.pl` script -- on each set of results. Following the recommendation of the DarkHorse documentation, we classified contigs as “positive for LGT ’’ if any ORF within the contig received a LPI score below 0.6.

#### WAAFLE vs. MetaCHIP

MetaCHIP^31^ is designed specifically to detect community-level LGT events among prokaryotes using a two-step process. The first step uses a homology-based approach to identify LGT via an all-against-all BLAST search of the provided input sequences or bins. The resulting best-match putative LGT candidates are then submitted to a phylogenetic validation check based on a set of 43 universal single-copy genes (SCGs)^61^ used to generate gene and species trees that are further reconciled using Ranger-DTL v2.0^62^. First, we attempted to assess MetaCHIP’s performance with the same synthetic contigs described above (i.e., those used to benchmark WAAFLE and DarkHorse). However, these input data would not progress beyond MetaCHIP’s phylogenetic validation step, likely due to its dependence on genomic data that are phylogenetically meaningful with a significant level of completeness^31^. Thus, we generated a compatible synthetic dataset to compare WAAFLE and MetaCHIP.

##### MetaCHIP-compatible synthetic contigs

To generate MetaCHIP-compatible synthetic contigs, we used a set of 20 human gut-native species previously used for bioBakery 2 testing (http://huttenhower.sph.harvard.edu/waafle) to construct contigs. Specifically, a random locus *R* in each recipient genome from this community was randomly selected and replaced by a randomly chosen locus *D* from a randomly selected donor genome, also from the community. This process was repeated 50 times to generate a total of 50 spiked LGT per each community genome. For each spiked community genome, we determined a set of breakpoints in the genome to generate synthetic contigs. A newly spiked LGT contig was generated by flanking each LGT locus with 1000 bp at both ends. Control (non-spiked) contigs were generated by shredding non-spiked regions into contigs of a defined length (by default, we used 2.5 kbp). Assuming that a set of contigs derived from a single community species (however complete) represents a discernible genome bin, synthetic bins were generated by sampling all the synthetic contigs from the same community genome.

We then ran MetaCHIP using default settings with the grouping rank *`-r`* option set to *`pcofgs`*. To further assess MetaCHIP’s dependence on bin completeness (and WAAFLE’s ambivalence), we ran both methods on the same inputs keeping 100%, 75%, 50%, and 25% of shredded control contigs per genome community.

### Analysis of HMP1-II metagenomes

#### Acquisition and assembly of sequencing data

We downloaded sequencing reads for 2,391 Human Microbiome Project (HMP1-II)^32,33^ metagenomes from http://hmpdacc.org. We then performed additional quality control on these reads using the KneadData workflow (v0.7.1, http://huttenhower.sph.harvard.edu/kneaddata). In brief, this workflow trims low-quality bases from sequencing reads, discards short reads, and additionally discards reads that aligned to the hg19 human reference genome (representing host contamination). We successfully assembled 2,376 of the downloaded metagenomes using MEGAHIT^63^ (v1.1.3) under default parameter settings. We discarded contigs <500 nt in length. These new metagenomic assemblies are available for download via http://huttenhower.sph.harvard.edu/waafle. Assembly statistics are provided in **Table S1**.

#### WAAFLE analysis

Each assembled metagenome was analyzed using the default WAAFLE workflow (as defined above). We used WAAFLE’s built-in gene calling feature (Step 1.5) to identify candidate protein-coding loci within contigs. We applied WAAFLE’s optional coverage-based quality control (Step 3) to exclude candidate LGT whose junctions neither had two or more supporting sequencing fragments nor consistent coverage relative to their flanking genes. After filtering out dubious LGT-containing contigs, we further discarded samples (assembled metagenomes) containing fewer than 1,000 total genes or which were flagged as ecological outliers in previous analyses of the HMP1-II dataset^32^. This left 2,003 assembled metagenomes for analysis. Unless otherwise specified, subsequent analyses are based on WAAFLE’s default outputs (describing the taxonomic and functional annotations of single-clade versus LGT-containing contigs), as detailed in the WAAFLE manual. These post-quality control output files are available for download from http://huttenhower.sph.harvard.edu/waafle.

#### Calculating LGT rates

We calculated rates of LGT for samples, clades, and clade pairs. In all cases, these rates take the form of “densities” of LGT events (i.e. discrete contigs identified by WAAFLE as containing an LGT) normalized by a background assembly size. This normalization accounts for the fact that all else being equal, we expect to see more LGT events as we assemble more contigs (from a sample or assigned to clades across samples). In addition, rates are characterized by their resolution, remoteness, and directedness, as defined in the main text. The LGT rate for a sample was defined as the number of LGT events in the sample divided by the total number of assembled genes from the sample (in 1,000s). For a given rate calculation, LGT events and background genes were required to meet a given taxonomic resolution (e.g., assignment to the genus level or lower), and LGTs were required to achieve a certain level of remoteness (e.g., an LCA for the transferring clades at the family-level or higher).

Rates of undirected LGT between a pair of clades (A, B) were defined as the number of A+B events seen across samples normalized to the total number of A and B genes assembled across samples (excluding repeated samples from the same individual and body site). Such rates can be interpreted as a robust average density of A+B transfers across metagenomic strains of A and B. Rates of directed LGT from B to A were similarly defined but normalized to the total number of A genes assembled across samples. Such rates can be interpreted as a robust average of the density of B-to-A transfers across metagenomic strains of A (i.e., the recipient clade in the transfer). Directed rates are systematically lower than their undirected counterparts due to the additional difficulty of identifying a directed LGT event with confidence. However, directed rates involving different donors and recipients are still comparable.

#### Computing phylogenetic distance and community abundance

We calculated the phylogenetic distance between two taxa using the ete3^64^ toolkit with the PhyloPhlAn reference tree^65^. If both taxa within an LGT pair were annotated to the species level (terminal nodes), distances were calculated by summing the branch length between them. If one or more taxa were annotated to a higher level, we 1) identified the internal node representing the last common ancestor (LCA) using all terminal nodes with the annotation (via regular expression), 2) calculated the average branch length between the LCA node and its terminal nodes, 3) calculated the branch length between the two taxon nodes (whether internal or terminal), and 4) summed distances from steps 2 and 3 for a final branch length. Relative abundances for community species (and higher level clades) in HMP1-II metagenomes were computed using MetaPhlAn2^56^ as previously described^32^.

#### Functional enrichment analysis

We performed two separate but related analyses of functional enrichment among novel LGT events in the human microbiome. The first and more precise analysis compared the frequencies of gene functional annotations in donated genes to all other genes across clades and samples from a given body site. This analysis thus focused on directed LGT events, where the LGT-donated genes could be distinguished from flanking genes of the recipient species. Because these events were ∼1/10th as common as their undirected counterparts (compromising coverage), we conducted a separate analysis of functional enrichments among genes found in LGT-containing contigs agnostic to the genes’ donor/recipient status. Thus, enrichment “signatures” from donated genes were diluted in this analysis by the inclusion of recipient genes in the “focal” gene set. In addition, if genes flanking LGT events are not a random sampling of background genes, then their corresponding functional enrichments will also be detected.

To conduct either functional enrichment analysis, we counted the number of total genes and LGT-associated genes with and without a given functional annotation across the assembled contigs from a given body site (ignoring repeated samples from the same individual). The significance of an annotation’s enrichment in the LGT-associated genes was then determined by Fisher’s exact test. The resulting two-tailed *p*-values were corrected within-body-site for multiple hypothesis testing using the Benjamini-Hochberg FDR method. We constrained biological interpretation to those functions that were 1) positively enriched (i.e., seen more often than expected in LGT-associated genes), 2) assigned to at least 10 LGT-associated genes in a single body site, and 3) whose enrichment was FDR significant at the *q*<0.1 level.

While the above methods are applicable to any collection of gene sets, we focused on Pfam^66^ domain annotations in this work. These annotations are advantageous in that they 1) are assigned objectively and with high coverage to proteins based on sequence homology, 2) could be inferred from WAAFLE’s existing UniRef90-level assignments to genes from metagenomic contigs, and 3) are reasonably well characterized. The mapping from UniRef90 to Pfam domains was extracted from HUMAnN2’s utility mapping databases (v0.11.1).

### Analysis of HMP2 stool metagenomes

#### Computational analysis and LGT selection

We further analyzed assembled metagenomes from control participants enrolled in the HMP2 Inflammatory Bowel Disease Multi’omics Database (IBDMDB) cohort^41^ (sometimes referred to as the IHMP or the Integrative Human Microbiome Project). Specifically, we applied the same read-level quality control and metagenomic assembly procedures introduced above in the context of HMP1-II metagenomes to the earliest-sampled stool metagenome from each non-IBD control participant. The resulting dataset included 26 assembled metagenomes from 26 unique individuals. We then applied WAAFLE to these samples, followed by an initial round of quality-control filtering as described above in the context of HMP1-II samples.

To further prioritize interesting LGT events for downstream experimental validation, we selected putative LGT-containing contigs that met the following conditions: 1) the contigs were gene-dense (>50% of nucleotide sites in a gene-coding locus); 2) the LGT junctions were not exceptionally long (<400 nt); 3) the transferring clades were resolved to at least the genus level, and 4) the clades’ donor and recipient roles could be inferred from gene adjacency. Of 616 initial LGT contigs, 27 passed both the general QC and supplemental prioritization filters. These 26 assembled HMP2 metagenomes and the subset of “interesting” LGT selected for validation are available from http://huttenhower.sph.harvard.edu/waafle. Additional data derived from and describing the IBDMDB cohort are available for download from http://ibdmdb.org.

#### PCR primer design and validation

We designed primers for LGT junctions from the above-selected putative LGT-containing contigs. As introduced above, an LGT junction is the transition between a gene from the recipient species and a gene from the donor species. It encompasses the 5’ and 3’ terminals of the respective genes and any intergenic space between them. Specifically, we 1) isolated 600-nt regions centered at each junction as target amplicons and then 2) ran Primer3^67^ (v2.4.0) with these regions as input using the following parameters:

● PRIMER_TASK=generic
● PRIMER_PRODUCT_SIZE_RANGE=250-350
● PRIMER_MIN_TM=48
● PRIMER_OPT_TM=50
● PRIMER_NUM_RETURN=20
● PRIMER_EXPLAIN_FLAG=1

We then manually selected primers from the returned list of candidates and ordered them from Life Technologies (**Table S8**).

Pre-extracted DNA from target HMP2 stool samples was retrieved from the Broad Institute repository. PCR was executed on a Bio-Rad T100 Thermocycler using the 5PRIME HotMasterMix (QuantaBio, catalog number 2200410). PCR cycling conditions were as follows: 1) 94°C for 3 min, 2) 94°C for 30 sec, 3) 48°C for 30 sec, 4) 72°C for 45 sec, 5) repeat steps 2-4 for 34 cycles (total 35 cycles), 6) 72°C for 10 min, and 7) 4°C hold. As a positive control, we amplified the 16S rRNA V4 region from a human oral swab provided by an anonymous donor using the 515F/806R primers from the Earth Microbiome Project (EMP)^68^. As negative controls, we performed PCR on two water samples using the same EMP primers. PCR products were checked on a 2% agarose gel containing 0.4 ug/ml of ethidium bromide in TAE buffer. They were run at 125 volts for 1 hr on an Owl A6 electrophoresis system (Thermo Scientific). PCR cleanup was performed using Agencourt Ampure XP Beads with 90 uL AMPure XP (Beckman Coulter, catalog number A63880).

## Data availability statement

HMP1-II metagenomes are available from the HMP DACC (http://hmpdacc.org) and from SRA BioProjects PRJNA48479 and PRJNA275349. Metadata and pre-computed taxonomic profiles for HMP1-II samples are also available from the HMP DACC. The HMP2 IBDMDB metagenomes used in this work are available from SRA BioProject PRJNA398089. Metadata for HMP2 samples are available from the IBDMDB website (https://ibdmdb.org). WAAFLE’s databases (as used in this work), synthetic evaluation contigs, HMP1-II and HMP2 assemblies, and LGT calls are available from the WAAFLE website (http://huttenhower.sph.harvard.edu/waafle).

## Code availability statement

WAAFLE is a free and open-source Python 3 software package available from GitHub (https://github.com/biobakery/waafle) and PyPI (https://pypi.org/project/waafle/). Installation and usage details, including links to download the databases and analysis products used in this work, are expanded on the WAAFLE website (http://huttenhower.sph.harvard.edu/waafle). Additional Python and R code used in the statistical analyses and visualizations from this work are available from the authors upon request.

## Inclusion and Ethics

The HMP1-II and HMP2 metagenomes and annotations used in this work are publicly available (**Data Availability Statement**). HMP2 stool samples from the Broad Institute were analyzed following the study’s per-institution Institutional Review Board approvals: overall Partners Data Coordination (IRB #2013P002215); MGH Adult cohort (IRB #2004P001067); MGH Pediatrics (IRB #2014P001115); Emory (IRB #IRB00071468); Cincinnati Children’s Hospital Medical Center (2013-7586); and Cedars-Sinai Medical Center (3358/CR00011696). The oral swab from an anonymous donor used as a positive control in the PCR validation experiment was not used for human subjects’ research purposes.

## Acknowledgments

This work was supported by the National Institutes of Health grants U54DE023798 (CH), R24DK110499 (CH), and K23DK125838 (LHN), the American Gastroenterological Association Research Foundation’s Research Scholars Award (LHN), and the Crohn’s and Colitis Foundation Career Development Award (LHN). The content is solely the responsibility of the authors and does not necessarily represent the official views of the NIH. We thank April Pawluk, Kelsey N. Thompson, Kristen Curry, and Todd Treangen for their comments on the manuscript and helpful discussions. We also acknowledge Monia Michaud, Casey Dulong, and Yan Yan for their help with validation experiments. Computational work was conducted on the FASRC Cannon cluster supported by the FAS Division of Science Research Computing Group at Harvard University.

## Supporting Information

### Supporting Figures

**Figure S1.**
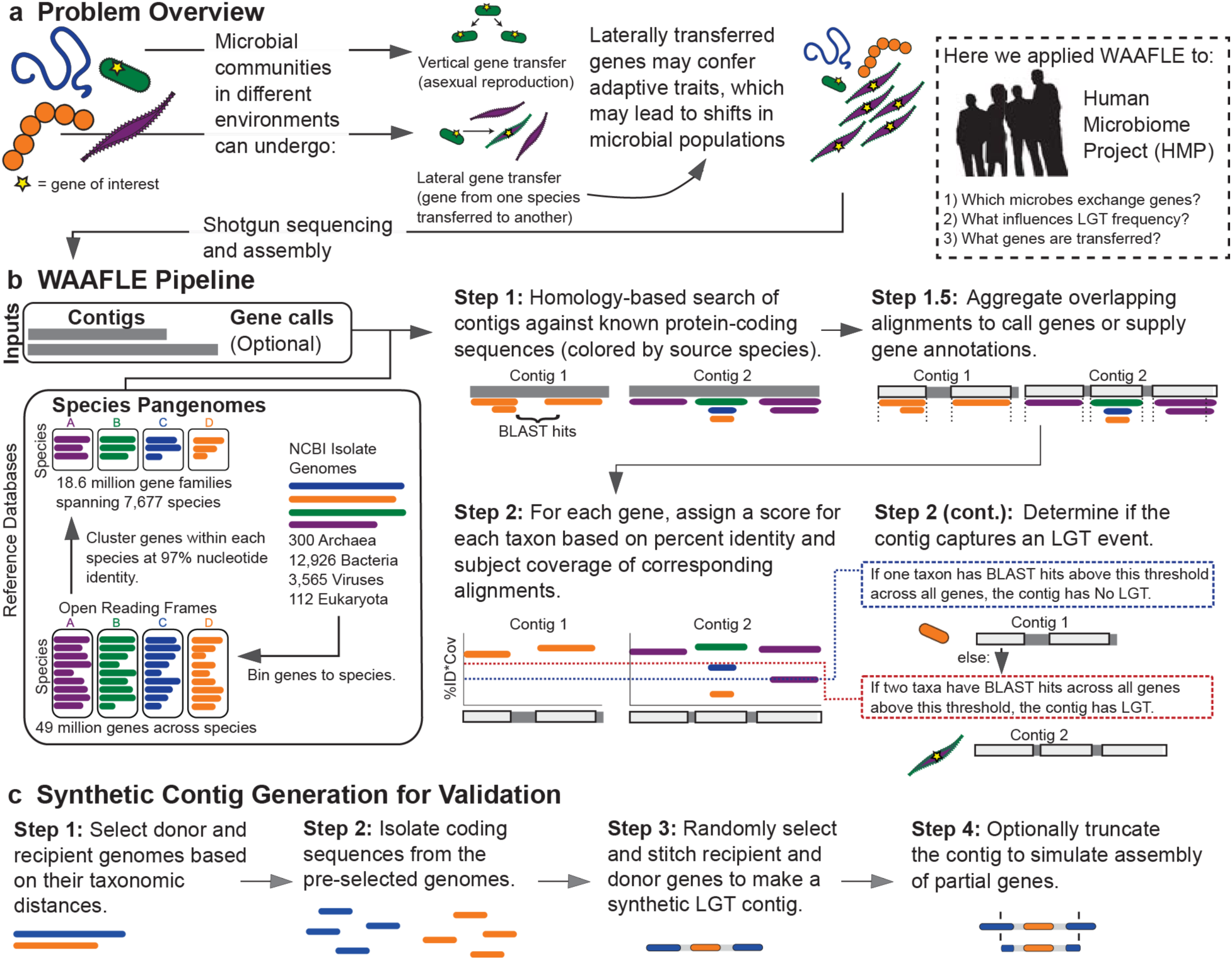
Details of the WAAFLE method and validation. (**A**) Within microbial populations, genes can be transferred vertically (from mother to daughter cell) or laterally (from a donor cell to a non-daughter recipient cell, often of another species). Lateral gene transfer (LGT) from a donor to a recipient may confer adaptive advantages to the recipient, leading to changes in community composition and function. (**B**) We designed WAAFLE to study LGT in microbial communities. WAAFLE identifies LGT events within assembled metagenomic contigs using a multi-step process. Step 1 of WAAFLE uses *blastn* to search contigs against a reference species pangenome database. This database was generated by downloading NCBI isolate genomes, binning isolate genes by species, and then clustering binned species genes at 97% nucleotide identity. WAAFLE can optionally call gene loci within contigs by clustering overlapping search results (Step 1.5). Step 2 of WAAFLE further analyzes search results to assign putative functions and per-taxon homology scores at each gene locus. If a single taxon has homology scores exceeding a threshold across all loci (k1, blue line), the contig is assigned to that taxon. If no taxon satisfies this criterion, WAAFLE then looks for pairs of taxa which collectively exceed a second, more stringent homology threshold at all loci (k2, red line). If a pair of taxa satisfy this criterion, WAAFLE predicts that the contig represents a LGT event between them. (**C**) To tune and evaluate WAAFLE, we generated LGT-containing synthetic contigs by selecting random donor and recipient genomes at different taxonomic levels and stitching their genes.

**Figure S2.**
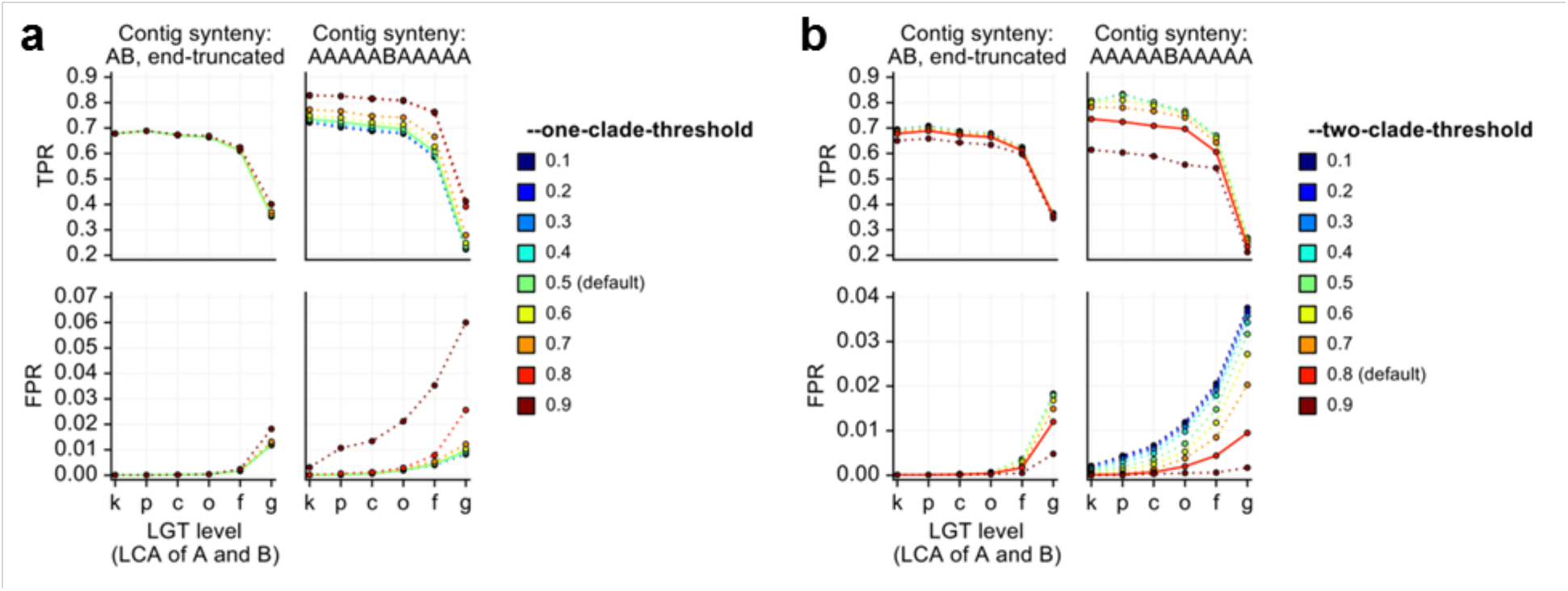
Tuning WAAFLE’s homology thresholds for identifying one- and two-clade contigs. **(A)** The “--one-clade-threshold” parameter, also referred to as k1, sets the minimum homology score that a single clade’s genes must meet in order to assign a contig to that clade. Higher values offer better sensitivity (TPR) but weaker specificity (FPR), especially for longer contigs. **(B)** The “--two-clade-threshold” parameter, also referred to as k2, sets the minimum homology score that a pair of clade’s genes must meet to assign a contig to that clade (assuming that no single clade has already exceeded “--one-clade-threshold”). Higher values offer better specificity but weaker sensitivity. WAAFLE’s default settings for these parameters are noted in the legend. These evaluations were conducted while holding out 20% of the database to simulate novel sequence data in synthetic contigs. LGT level, i.e. the LCA of the two species participating in the LGT, is given as the abbreviated taxonomic rank of the LCA (“k” for “kingdom”, “p” for “phylum”, and so forth).

**Figure S3.**
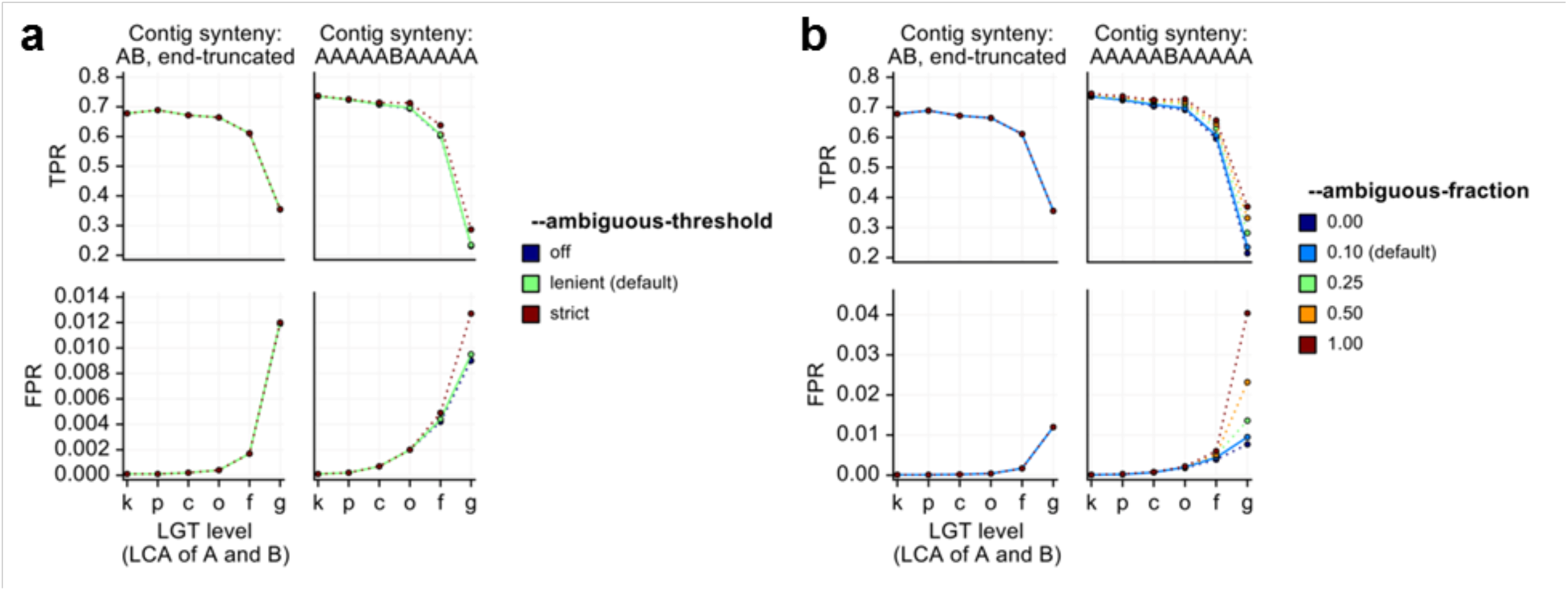
Tuning WAAFLE’s thresholds for excluding LGT with large fractions of ambiguous loci. **(A)** The “--ambiguous-threshold” parameter sets the allowed threshold for calling a particular locus “ambiguous” in an A+B putative LGT event. “Off” means that the locus will be flagged as ambiguous if A and B each have a hit to the locus (regardless of homology score), lenient requires both scores to exceed k1=0.5, and strict requires both scores to exceed k2=0.8. Stricter values had minimal impact on accuracy. **(B)** The “--ambiguous-fraction” parameter sets the maximum allowed fraction of sequence length contributed by ambiguous loci. Higher values result in considerably reduced specificity at the intragenus level. WAAFLE’s default settings for these parameters are noted in the legend. These evaluations were conducted while holding out 20% of the database to simulate novel sequence data in synthetic contigs. LGT level, i.e. the LCA of the two species participating in the LGT, is given as the abbreviated taxonomic rank of the LCA (“k” for “kingdom”, “p” for “phylum”, and so forth).

**Figure S4.**
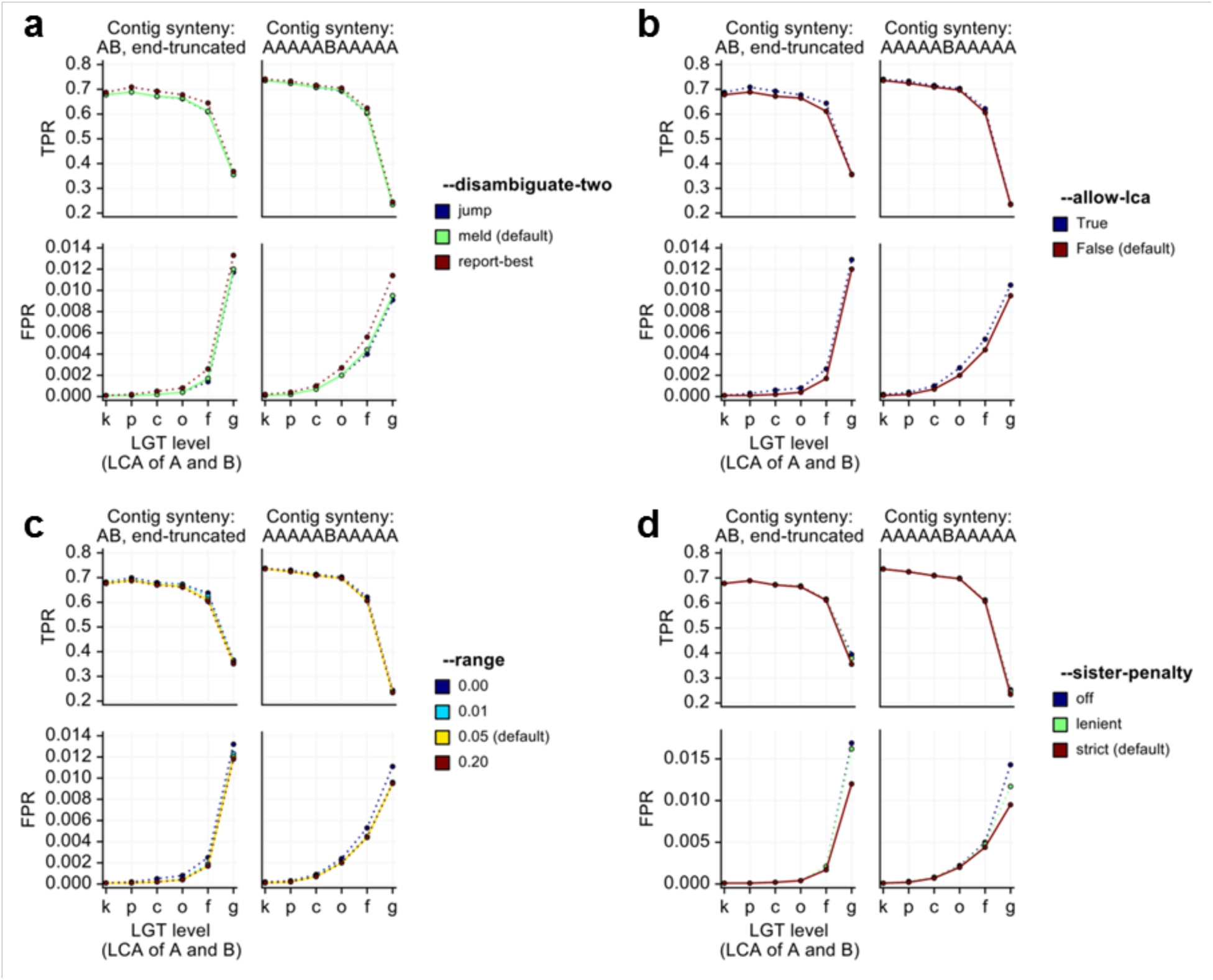
Tuning other notable parameters of the WAAFLE algorithm. **(A)** The “--disambiguate-two” parameter sets WAAFLE’s behavior when multiple LGT-based explanations are found for a contig. “Jump” reevaluates the contig at the next level up in the taxonomy, “meld” reduces putative recipients and donors to their respective LCAs, and “report-best” reports the single explanation with the best average per-locus score. “Report-best” suffers from slightly reduced specificity at the intragenus level. **(B)** The “--allow-lca” parameter determines whether or not WAAFLE will report an A:B LGT where A is also the LCA of A and B (e.g. a transfer of the form “*Bacteroides vulgatus* to *Bacteroides* unclassified”). Such cases tended to be rare but were slightly enriched for false positive calls. **(C)** The “--range” parameter determines the extent of alternative LGT solutions considered below the best-scoring explanation. Using a non-zero value (i.e. considering alternative explanations that were close to the best explanation) offered slightly better specificity. **(D)** The “--sister-penalty” applies to A:B LGTs in which at least one B locus also scored well in a sister clade of A. The “off” setting does not apply the penalty, the “lenient” setting requires the sister homolog to exceed k2, and the “strict” setting requires the sister homolog to only exceed k1. Stricter settings offered improved specificity at the intragenus level with essentially no cost in sensitivity. WAAFLE’s default settings for these parameters are noted in the legend. These evaluations were conducted while holding out 20% of the database to simulate novel sequence data in synthetic contigs. LGT level, i.e. the LCA of the two species participating in the LGT, is given as the abbreviated taxonomic rank of the LCA (“k” for “kingdom”, “p” for “phylum”, and so forth).

**Figure S5.**
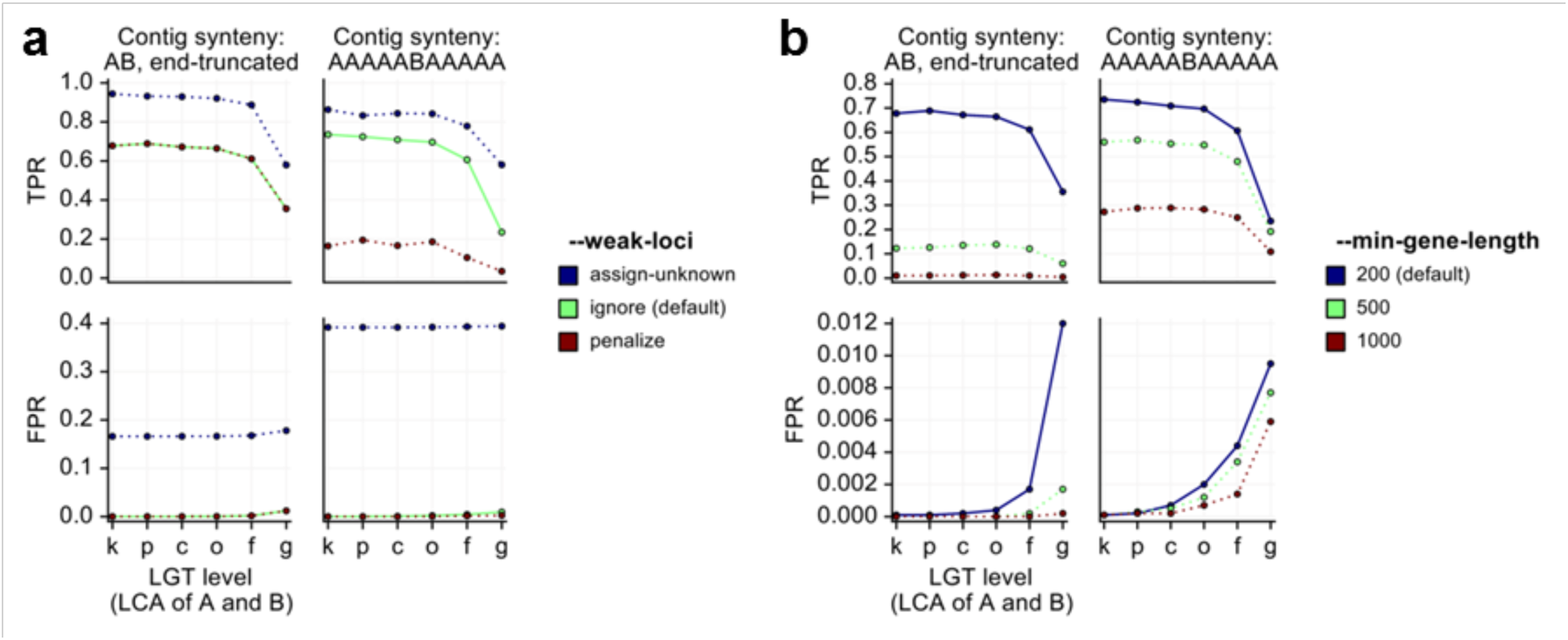
Tuning WAAFLE’s parameters related to identification and exclusion of low-confidence gene loci. **(A)** The “--weak-loci” parameter determines WAAFLE’s handling of a locus that never exceeded the k1 (lower) homology threshold in any species. The “assign-unknown” setting introduces an unknown species with score 1 - max(observed scores), which dramatically reduces specificity. The “penalize” setting does not treat these loci differently from others, and results in dramatically reduced sensitivity. The “ignore” setting removes these loci from the clade-by-locus homology scoring table and offers good sensitivity and excellent specificity. **(B)** The “--min-gene-length” parameter filters short loci from the clade-by-locus homology scoring table. Larger values result in dramatic loss of sensitivity with only minor gains in specificity. WAAFLE’s default settings for these parameters are noted in the legend. These evaluations were conducted while holding out 20% of the database to simulate novel sequence data in synthetic contigs. LGT level, i.e. the LCA of the two species participating in the LGT, is given as the abbreviated taxonomic rank of the LCA (“k” for “kingdom”, “p” for “phylum”, and so forth).

**Figure S6.**
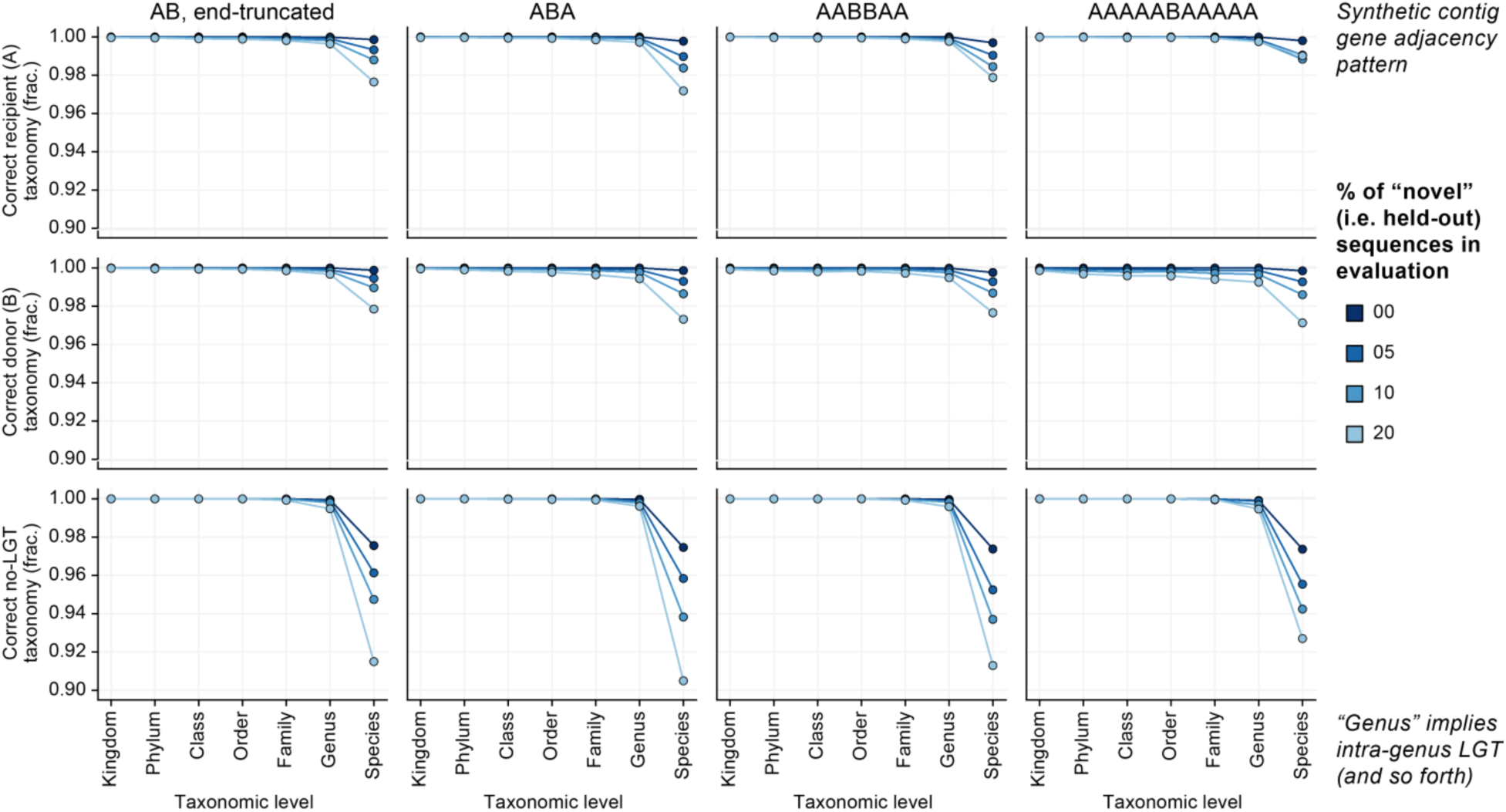
Accuracy of WAAFLE’s taxonomic assignments for correctly called LGT and non-LGT contigs. The top row reports the accuracy of WAAFLE’s call for the recipient species in an LGT contig, the middle row reports the accuracy for the donor species, and the bottom row reports the accuracy of the single species in non-LGT (negative control) contigs. In all cases, and for a range of gene adjacency/gene order configurations, accuracy is very high for all taxonomic levels (>96%) when using the full pangenome database for evaluation. Holding out fractions of this database (up to 20% of each species pangenome) tends to reduce the accuracy of species-level calls (though never below 90%), while genus-level calls and higher remain very accurate (>99%).

**Figure S7.**
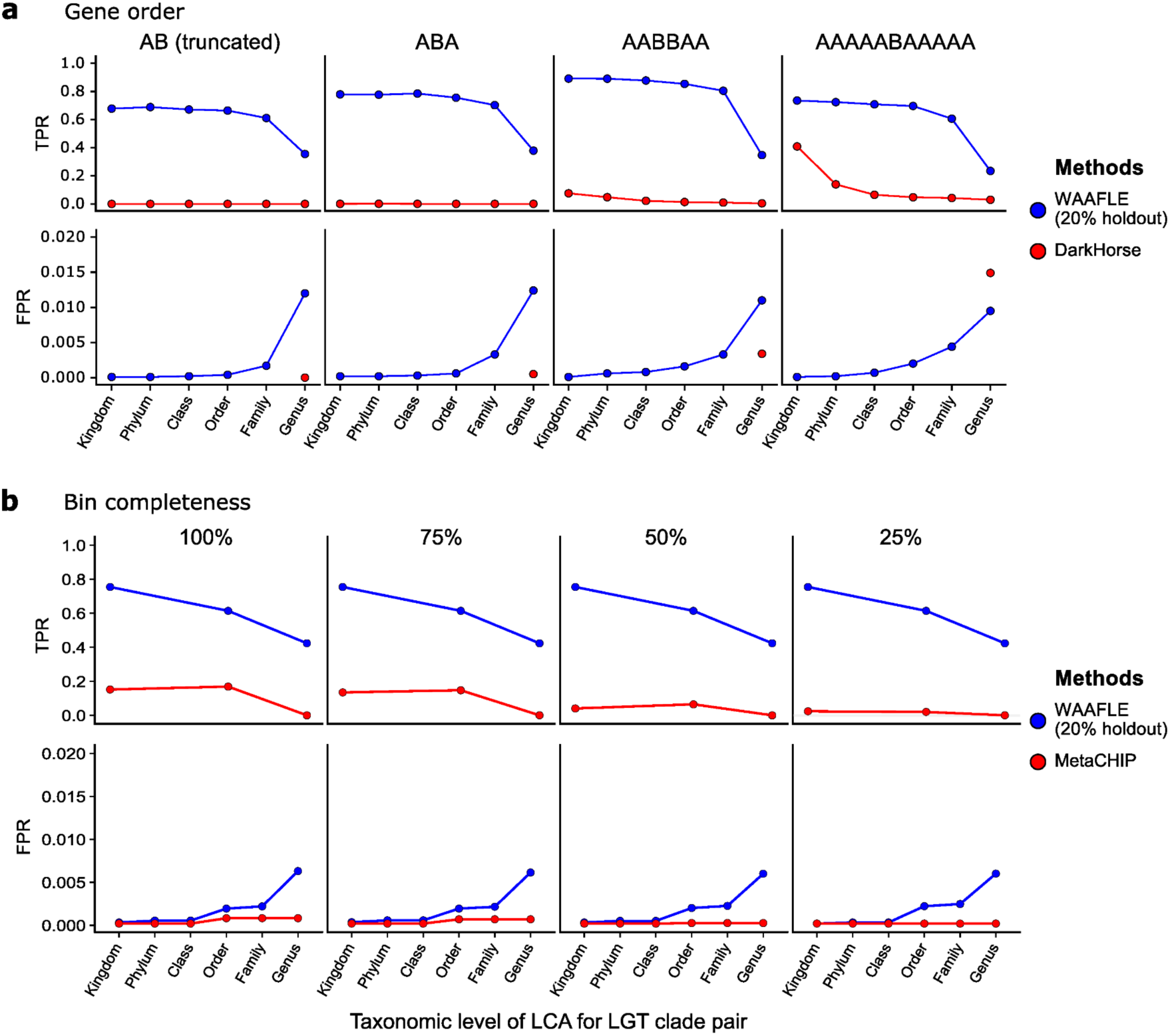
Relative accuracy of WAAFLE, DarkHorse, and MetaCHIP on synthetic LGT and control contigs. WAAFLE was penalized with a 20% holdout of its search database, while DarkHorse was evaluated using a translated version of the complete database, and MetaCHIP was evaluated without further constraints on its respective input. (**A**) DarkHorse only achieved non-negligible sensitivity (TPR) for the longest contigs (rightmost column) containing the most “extreme” LGT events (i.e. between pairs of species with the kingdom- or phylum-level LCAs). WAAFLE’s specificity (FPR) is stratified according to the taxonomic level of the LGT LCA as in Fig. 1 from the main text (e.g. an intra-genus false positive is counted as a true negative at the family level; x-axis). This level of stratification was not possible for DarkHorse, and so a single FPR value is plotted at “genus” resolution for comparison. DarkHorse offered better specificity than WAAFLE on shorter contigs (where it made relatively few LGT calls) but not on the longest contigs. (**B**) Here, an additional comparison was performed between WAAFLE and MetaCHIP using a separate synthetic dataset designed for MetaCHIP compatibility. TPR and FPR were computed and plotted as in ‘(A)’ with TPR calculations restricted to taxonomic ranks assigned to at least 100 LGT LCAs (i.e. kingdom, order, and genus). Results are stratified according to the completeness of the metagenomic bins into which LGT and control contigs were grouped. WAAFLE’s sensitivity here was similar to that observed in the preceding evaluations and consistently higher than MetaCHIP. While MetaCHIP’s specificity was correspondingly very high, WAAFLE again exhibited a peak FPR of only ∼0.5% at the intra-genus level, improved at higher ranks. Notably, WAAFLE’s performance was not dependent on bin completeness, while MetaCHIP proved less sensitive to LGT events in less-complete bins.

**Figure S8.**
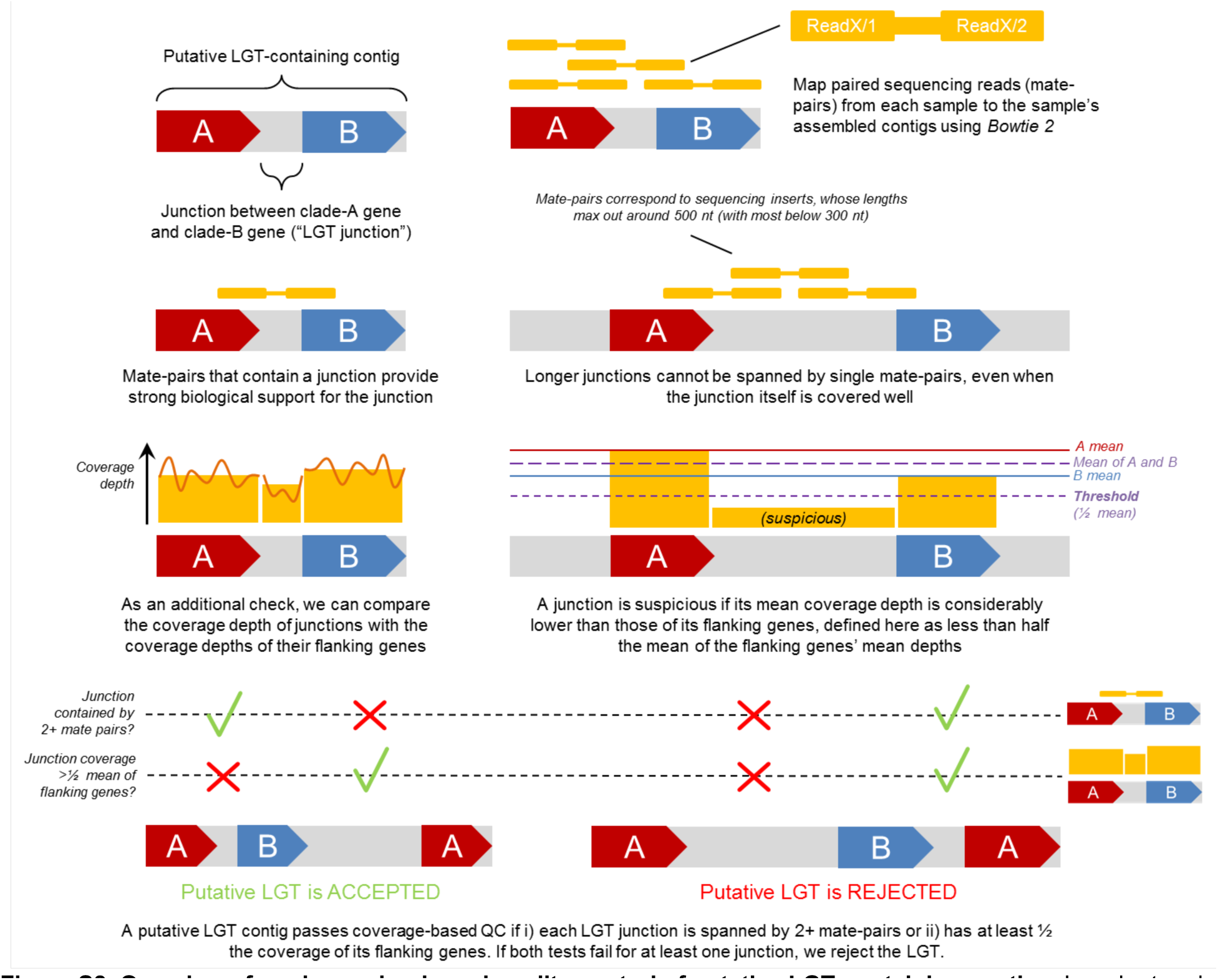
Overview of read-mapping-based quality control of putative LGT-containing contigs. In order to rule out potential misassembly events that appear as LGT events, we mapped sequencing reads from each metagenome to the metagenome’s assembled contigs. We then required each LGT junction in putative LGT contigs to either i) be spanned by individual mate-pairs or ii) have reasonable coverage relative to the coverage of the flanking genes. The former criterion is more applicable for shorter junctions, while the latter criterion is more applicable for longer junctions.

**Figure S9.**
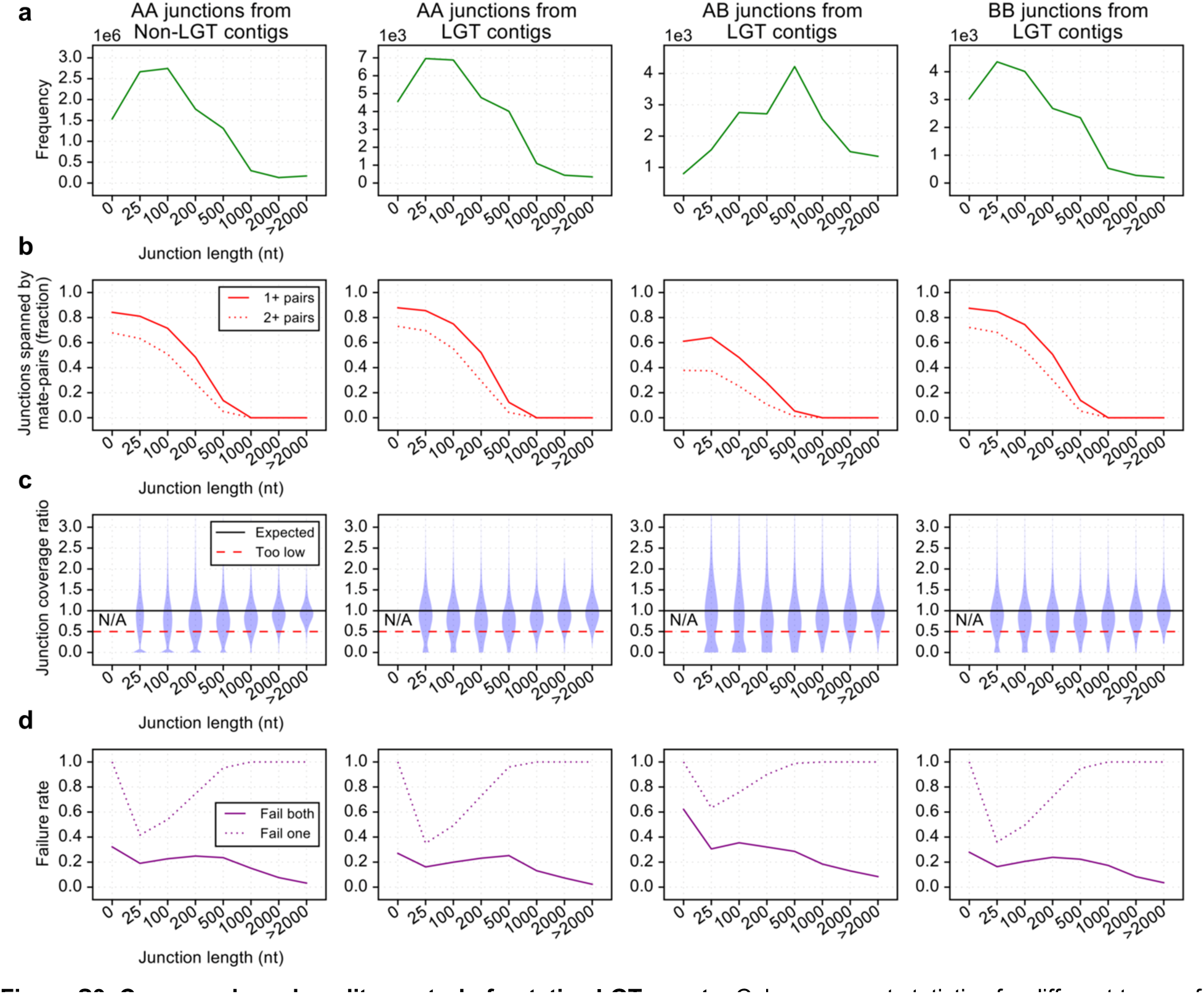
Coverage-based quality control of putative LGT events. Columns report statistics for different types of gene-gene junctions across non-LGT and LGT contigs. AB junctions from LGT contigs (i.e., junctions between genes from different clades) were required to pass quality filters for their source contigs to be included in downstream analyses. **(A)** Frequencies of different junction types stratified by length. Length 0 indicates that the second gene in the pair starts at or before the end of the first gene. **(B)** Fractions of junctions contained within a sequencing fragment (mate-pair). Junctions longer than typical sequencing insert sizes receive less support here by definition. **(C)** Ratios of coverage depth in the gene-gene junction to the average coverage depth of the two flanking genes. This value cannot be computed for junctions in the “0 nt” category and is less reliable for shorter junctions. **(D)** A junction was considered to pass coverage-based quality control if it passed one of the following two tests: i) the junction was spanned by at least two mate-pairs, or ii) the junction’s coverage ratio exceeded 0.5. Because mate-pair containment is generally not feasible for longer junctions, and coverage ratios are less reliable for shorter junctions, requiring junctions to pass both of these filters would exclude the vast majority of junctions.

**Figure S10.**
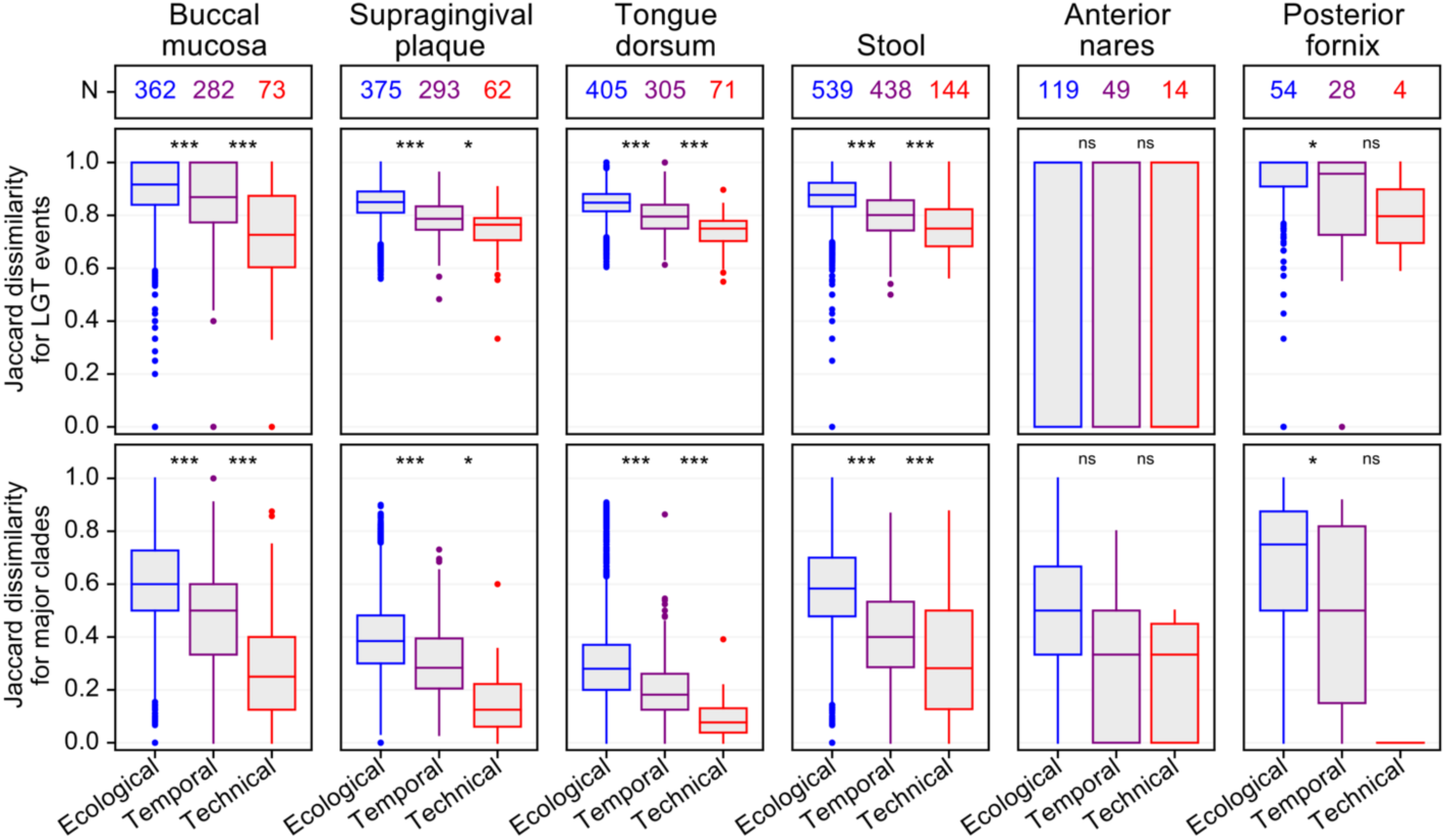
Relative similarity of HMP1-II sample LGT profiles and background taxonomy from metagenomic assembly. Top row: Numbers of unique HMP1-II participants contributing similarity values to the boxplots below. Middle row: Jaccard similarity of inter-genus LGT profiles. “Ecological” encompasses comparisons between samples from different participants. “Temporal” encompasses comparisons between longitudinal samples from the same subject. “Technical” encompasses comparisons between repeated samples from the same biological specimen (unique to a person, body site, and time point). Bottom row: Comparisons between the sets of “major” genera detected from sample assembly (500+ genes assembled). Distributions were compared by Wilcoxon Rank-Sum tests with two-tailed *p*-values; “***” implies *p*<0.001, “**” implies *p*<0.01, “*” implies *p*<0.05, and “ns” implies *p*≥0.05.

**Figure S11.**
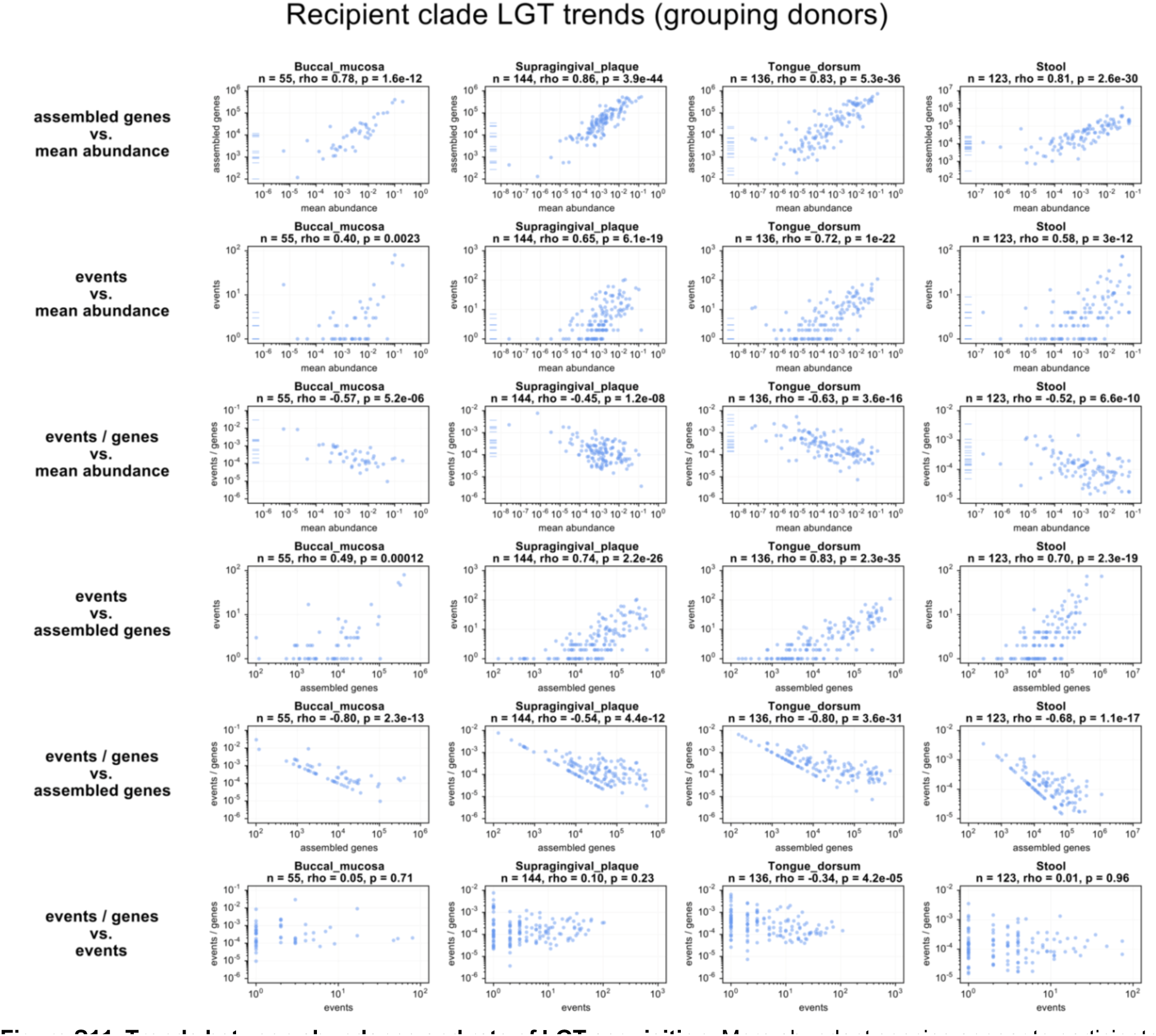
Trends between abundance and rate of LGT acquisition. More abundant species appear to participate in more LGT events as a recipient due in part to the confounding of abundance and assemble-ability. Density of events as an LGT recipient (“events / genes”) appears to negatively correlate with abundance. However, this trend is dominated by the low LGT densities of low-abundance species, where density varies as ∼1/(assembly size), thus inducing a negative correlation with abundance.

**Figure S12.**
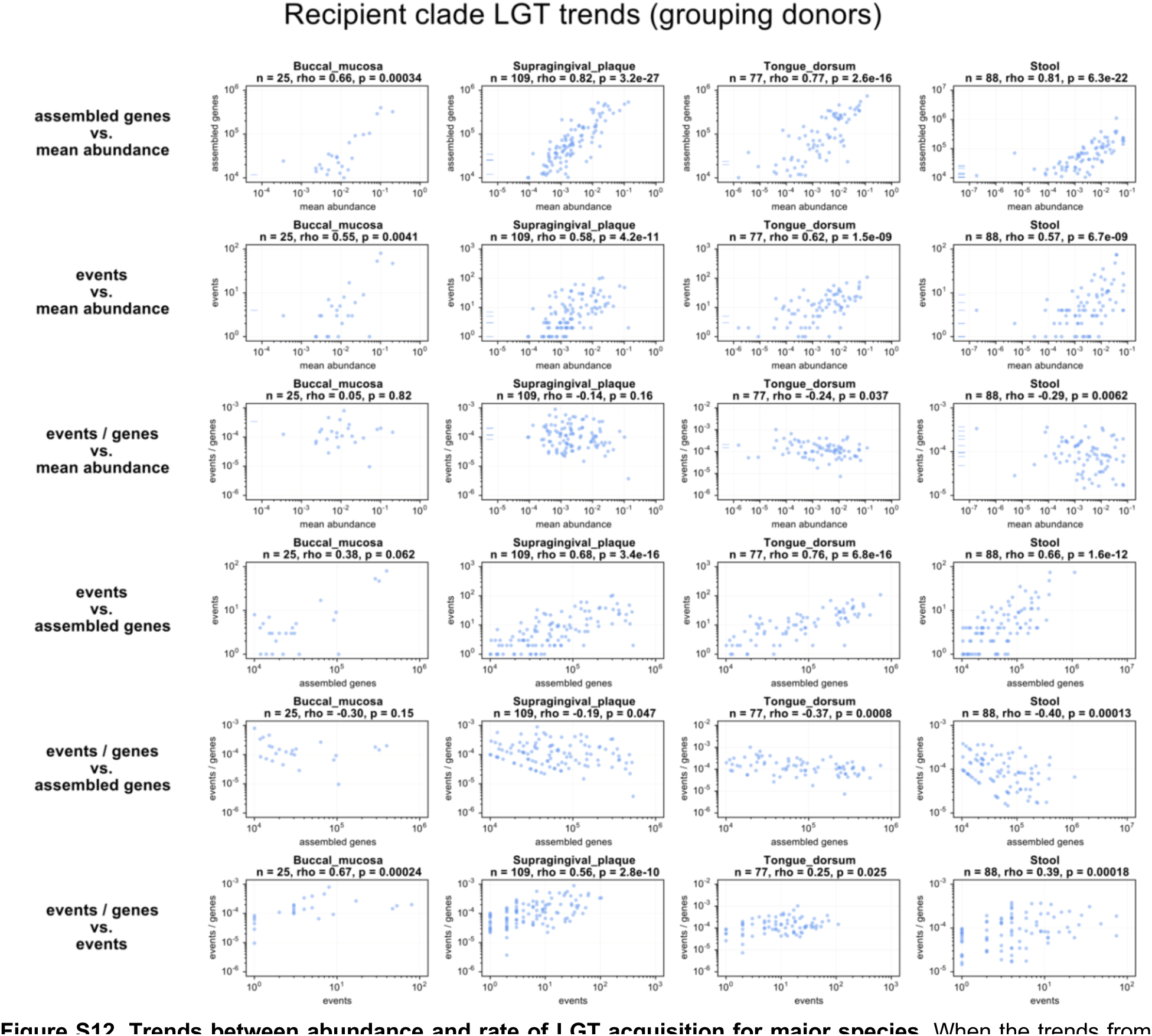
Trends between abundance and rate of LGT acquisition for major species. When the trends from **Fig. S11** are re-investigated among major species only (>10K assembled genes across samples), the spurious negative correlation between density of events as an LGT recipient (“events / genes”) and abundance becomes much less pronounced, and no significant positive correlation is observed.

**Figure S13.**
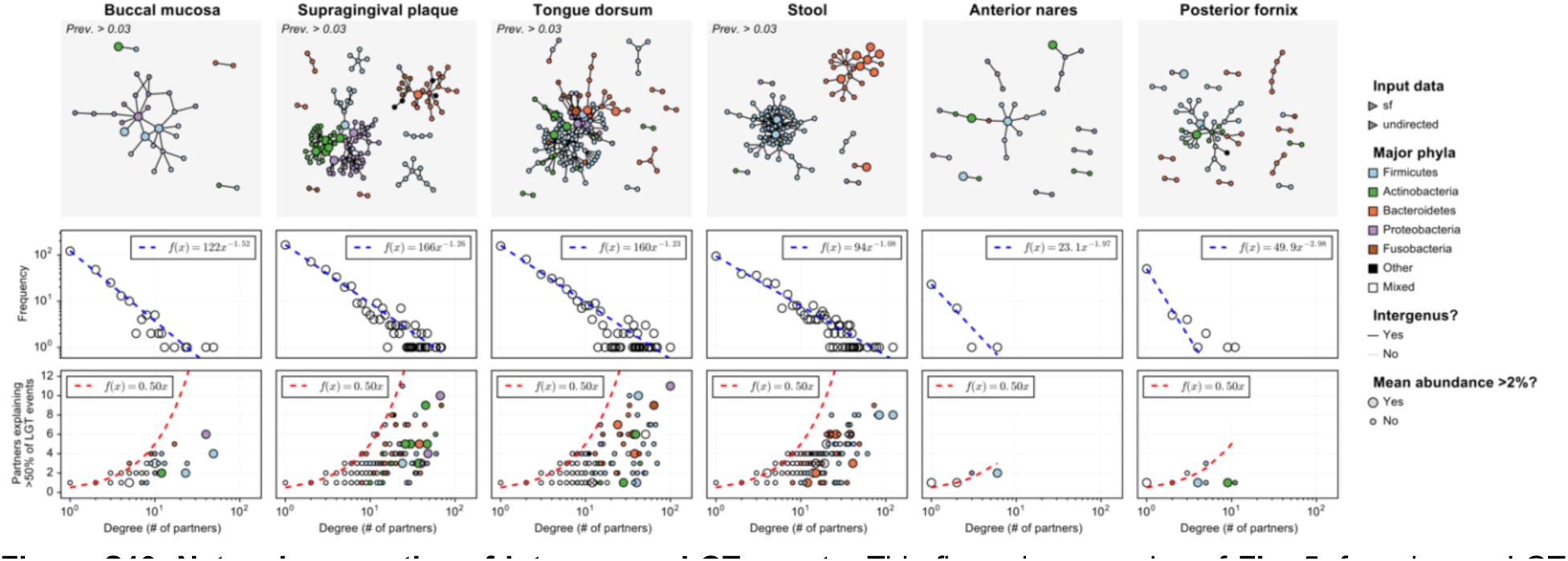
Network properties of inter-genus LGT events. This figure is an analog of Fig. 5, focusing on LGT events resolved to the genus level with LCA at or above the family level.

**Figure S14.**
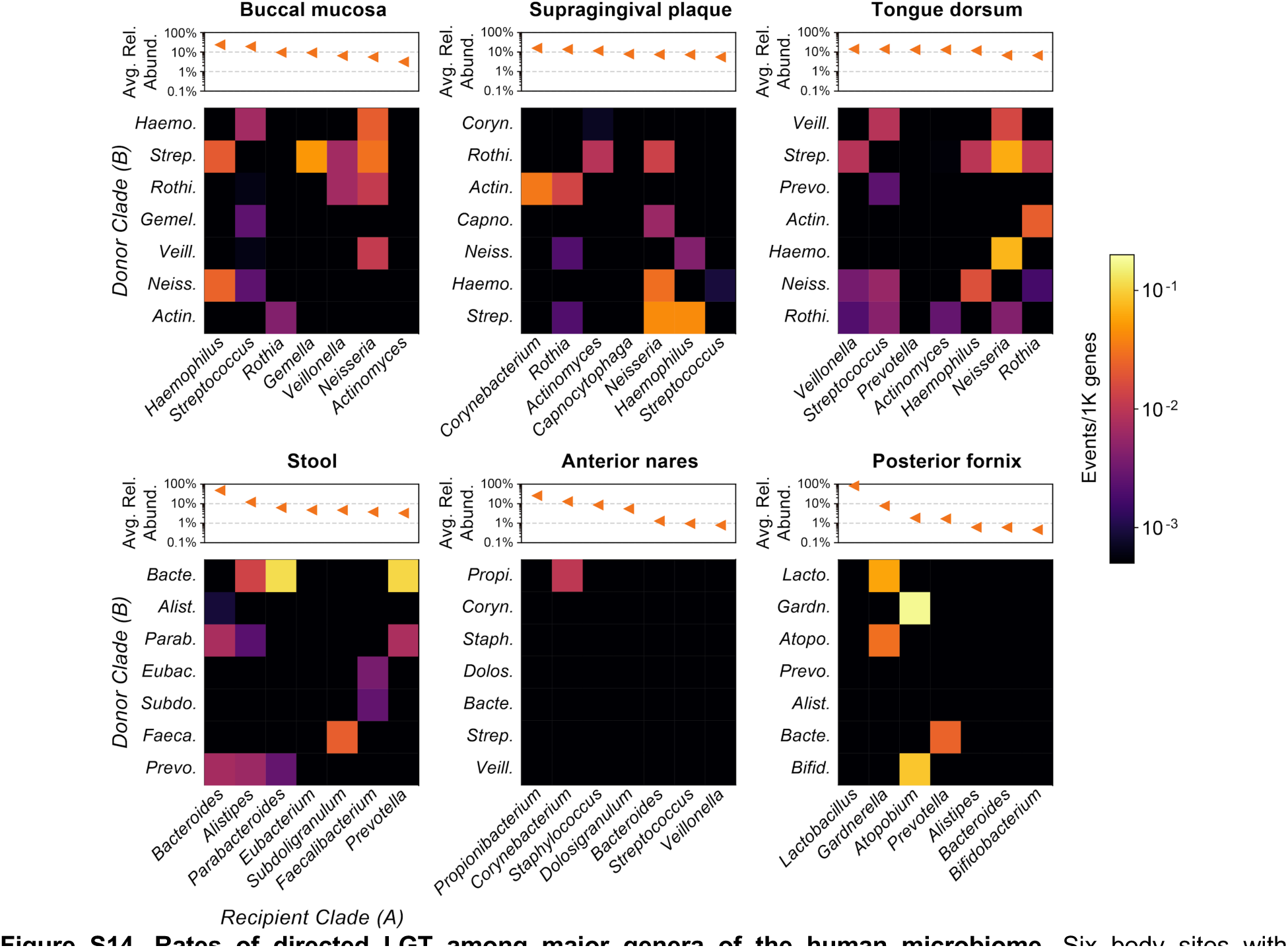
Rates of directed LGT among major genera of the human microbiome. Six body sites with metagenome sequencing from at least 20 individuals are shown; the three body sites in the top row are all from the oral cavity. Heatmap values indicate the rate of undirected LGT between major genera from the body site, with “major genera” defined based on ranked average relative abundance. Rates are computed over first-visit samples from HMP1-II participants.

**Figure S15.**
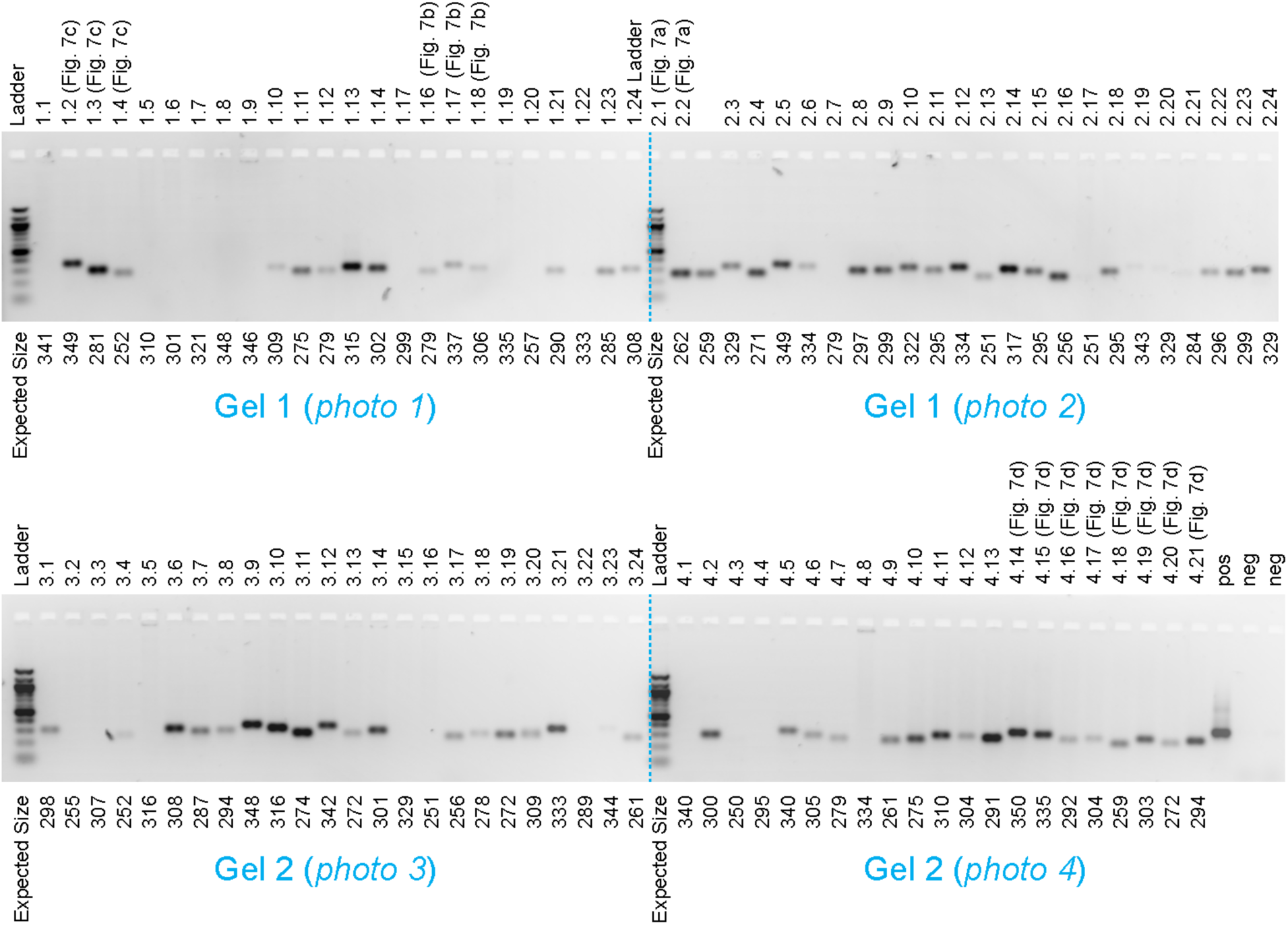
Gel images for PCR of LGT junctions. Reactions referenced in Fig. 7 **A**-**D** are highlighted; all reactions are numbered (x.y) for cross-reference to **Table S8**. Expected sizes for PCR products are labeled underneath lanes. Positive control (“pos”) is an amplification of 16S rRNA V4 region from a human oral swab; negative controls (“neg”) are reactions of water using 16S primers (**Methods**). “Ladder” is a New England BioLabs 100 bp ladder (catalog #N3231). Dashed blue lines indicate digital stitching over separate photographs.

**Figure S16.**
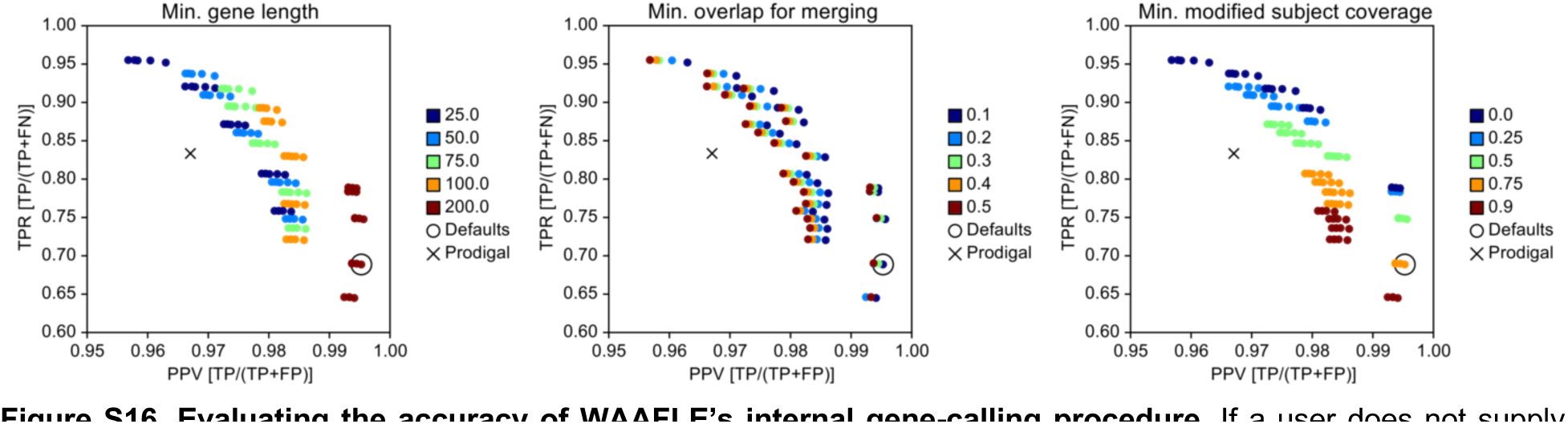
Evaluating the accuracy of WAAFLE’s internal gene-calling procedure. If a user does not supply WAAFLE with independently generated ORF calls for their input contigs, WAAFLE can approximate the locations of gene-coding loci within those contigs based on the results of its homology-based search step. Here, evaluated on random contigs drawn from sequenced isolate genomes (including partial genes), WAAFLE was able to recover a majority of known genes while producing small numbers of false positive calls (similar to the dedicated ORF caller Prodigal). WAAFLE’s default settings (circled) are less sensitive but more specific than Prodigal due to the exclusion of shorter candidate genes (<200 nt). Such genes were more likely to suggest false positive LGT events in downstream steps, and so they are excluded by default during gene calling.

### Supporting Tables (captions)

**Table S1. Assembly statistics for 2,384 HMP1-II metagenomes.** Metagenomes were assembled with MEGAHIT v1.1.3. Assembly statistics (e.g. N50 score) were computed using assembly-stats v1.0.1 (cloned from http://github.com/sanger-pathogens/assembly-stats). Contigs <500 nt were discarded before calculating statistics.

**Table S2. Counts and rates of LGT events from HMP1-II metagenomes.** Columns indicate body site, clade pair, clade taxonomy, number of LGT events, and LGT rate (events normalized to clade pair assembly size).

**Table S3. Counts and rates of inter-genus LGT events from HMP1-II metagenomes.** Columns indicate focal body site, clade pair, clade taxonomy, number of LGT events, and LGT rate (events normalized to clade pair assembly size). Events between species in the same genus are excluded.

**Table S4. Counts and rates of directed LGT events from HMP1-II metagenomes.** Columns indicate focal body site, donor/recipient clades, donor/recipient taxonomy, number of LGT events, and LGT rate (events normalized to recipient assembly size).

**Table S5. Counts and rates of directed inter-genus LGT events from HMP-II metagenomes.** Columns indicate focal body site, donor/recipient clades, donor/recipient taxonomy, number of LGT events, and LGT rate (events normalized to recipient assembly size). Events between species in the same genus are excluded.

**Table S6. Molecular functions (Pfam domains) enriched in transferred genes across body sites relative to background genes.** Table fields indicate the body site, Pfam ID, Pfam name, overlap (instances of the domain in transferred genes), expected overlap (assuming that transferred domains are a random sampling of all domains), fold enrichment (the ratio of the overlap to the expected overlap), two-tailed *p*-value from Fisher’s exact test, and Benjamini-Hochberg FDR *q*-value (computed within-site).

**Table S7. Molecular functions (Pfam domains) enriched in LGT-containing contigs across body sites relative to background contigs.** Table fields indicate the body site, Pfam ID, Pfam name, overlap (instances of the domain in LGT contigs), expected overlap (assuming that LGT contig domains are a random sampling of all domains), fold enrichment (the ratio of the overlap to the expected overlap), two-tailed *p*-value from Fisher’s exact test, and Benjamini-Hochberg FDR *q*-value (computed within-site).

**Table S8. PCR primers and results from the LGT junction amplification experiment.** Each row corresponds to a primer pair and PCR reaction targeting one of the two junctions from a putative LGT event. Sample IDs refer to control stool metagenomes from ibdmdb.org. Longer junctions were divided into >1 tiled segment (as indicated in the “Segment” column). The “Gel Label” column associates rows of this table with the gel images in Fig. 7 and **Fig. S15**. Quantification values were determined from a Qubit fluorometer.

